# The response of desert biocrust bacterial communities to hydration-desiccation cycles

**DOI:** 10.1101/2021.04.04.438350

**Authors:** Capucine Baubin, Noya Ran, Hagar Siebner, Osnat Gillor

**Author notes:** Corresponding authors: Capucine Baubin, Zuckerberg Institute for Water Research, Blaustein Institutes for Desert Research, Ben-Gurion University of the Negev, 8499000, Israel., Osnat Gillor, Zuckerberg Institute for Water Research, Blaustein Institutes for Desert Research, Ben-Gurion University of the Negev, 8499000, Israel.

## Abstract

Rain events in arid environments are highly unpredictable, interspersing extended periods of drought. Therefore, following changes in desert soil bacterial communities during hydration-desiccation cycles in the field, was seldom attempted. Here, we assessed rain-mediated dynamics of active community in the Negev Desert biological soil crust (biocrust), and evaluated the changes in bacterial composition, potential function, and photosynthetic activity. We predicted that increased biocrust moisture would resuscitate the phototrophs, while desiccation would inhibit their activity. Our results show that hydration increased chlorophyll content, resuscitated the biocrust *Cyanobacteria,* and induced potential phototrophic functions. However, decrease in the soil water content did not immediately decrease the phototrophs activity, though chlorophyll levels decreased. Moreover, while the *Cyanobacteria* relative abundance significantly increased, *Actinobacteria*, the former dominant taxa, significantly decreased in abundance. We propose that, following a rain event biocrust moisture significantly decreased, almost to drought levels, yet the response of the active bacterial community lagged, in contrast to topsoil. Possible explanations to the described rain-mediated bacteria dynamics are discussed.

## 1. INTRODUCTION

Arid environments are the largest terrestrial biomes on Earth and accounts for 35% of the landmass (Pointing and Belnap, 2012). Since rain in arid environments is rare and unpredictable, the main source of water is dew (Malek et al., 1999) and fog (Kidron et al., 2002). This moisture is readily absorbed to the soil surface, but would quickly evaporates due to high temperatures and low humidity (Cameron and Blank, 1966). The long droughts in drylands limit plant growth and in their stead, the soil is covered by microbial mats, called biological soil crust (biocrust). Biocrusts are a matrix of phototroph and heterotroph microorganisms that bind together with soil particles, by using extracellular polymeric substances (EPS) (Belnap and Lange, 2001; Campbell et al., 1989; Kidron et al., 2020).

Biocrust phototrophs are the main primary producers in this desolate habitat and together with the heterotrophs form a rigid and stable mat that is able to resist to xerification and soil erosion (Aanderud et al., 2019; Bowker et al., 2018). Biocrusts are the main source of carbon and nitrogen (Agarwal et al., 2014), and a strong contributor of soil respiration (Castillo-Monroy et al., 2011) in deserts. It was recently shown that, during long droughts many of the biocrust microorganisms rely not only on photosynthesis but also on oxidation of atmospheric trace gases (Leung et al., 2020; Meier et al., 2021). Yet, once the biocrust is hydrated, the phototrophs respond quickly by inducing their photosynthetic systems and related functions to take full advantage of the rare water abundance before the soil dehydrates (Murik et al., 2017). To that end, photosynthetic members of the biocrust community form a seed bank of species that are able to spring to life whenever the water content increases (Kedem et al., 2020; Lennon and Jones, 2011; Murik et al., 2017). Yet, the abrupt hydration may also cause osmotic shock that could result in massive cell lysis and the release of osmoregulatory solutes (Halverson et al., 2000; Harris, 1981). The period of water abundance is usually brief, and the soil quickly dehydrates forcing the bacteria to cease their activity (Murik et al., 2017; Oren et al., 2019). Therefore, the members of the biocrust community must respond quickly and efficiently not only to hydration but also to the subsequent desiccation.

Earlier studies focused on community structure and cyanobacterial response to hydration-desiccation cycles under controlled conditions (Angel and Conrad, 2013; Meier et al., 2020; Oren et al., 2019; Wu et al., 2013). To the best of our knowledge, these cycles were never monitored in the field during a rain event. Under natural conditions, the biocrust community dynamics of the hydration-desiccation cycle may be affected by plethora of variables, such as temperature, rain intensity, or soil local structure, which could not be applied in a laboratory setting. Thus, it is imperative to elucidate the resuscitated community and its response to the gradual dehydration after a rain event in the field.

In this study, we followed the community structure and activity before, during, and after a rain event in the arid Negev Desert (Israel). We studied the active biocrust community by using SSU ribosomes as a proxy to active bacterial community (Št’ovíček et al., 2017). Although ribosomes do not quickly degrade in dormant or even dead cells (Sukenik et al., 2012; Sunyer-Figueres et al., 2018), under field conditions they present a reliable mean to distinguish between active and inactive cells (Angel et al., 2013; Št’ovíček et al., 2017). We hypothesised that the biocrust community would quickly respond to hydration and to desiccation. We predicted that high soil moisture would trigger photosynthetic activity and a decreasing soil moisture will lead to an inactivation of the phototrophs within the biocrust community. We further predicted that heterotrophs response to hydration-desiccation would differ among phyla as previously reported for biocrust (Angel and Conrad, 2013) and topsoil (Št’ovíček et al., 2017) collected from the same site. The most apparent change detected in both soil horizons was the sharp decrease in the relative abundance of *Actinobacteria* that dominant the soil during droughts but decline upon hydration.

## 2. MATERIAL AND METHODS

### 2.1. Sampling

The study was conducted in the long-term ecological research station in the Negev Desert Highlands (Zin Plateau, 30°86’N, 34°80’E, Israel; Figure 1). In this arid environment, the average annual rainfall is around 90 mm and extends from October to April. Biocrust samples were collected on 20/06/17 during the dry season (T[0]; average temperature: 32.4°C) and during a rain event in the wet season from 29/01/18 through 01/02/18 at 24 hr intervals. The rain event (5.1 mm, maximum average temperature 14.6 °C) occurred 29/01/18 (T[R]) and samples were collected till the biocrust dried (T[1], T[2], T[3]; Figure 1) For each time point, five samples at least 10 m apart were collected (N = 25 samples). The biocrust samples were homogenised using a 2 mm sieve and then four subsamples were stored: (1) at −80°C for molecular analysis; (2) at −20°C for chlorophyll extraction; (3) at 60°C for 3 days and then kept at room temperature for chemical analysis; and (4) was used immediately to evaluate the water content.

### 2.2. Physico-chemical analyses

Water content, organic carbon and total nitrogen were measured in the soil samples. Biocrust water content was determined by the gravimetric method, the soil was weighed before and after oven drying at 105 C, then the percentage of moisture in the soil was determined (Scrimgeour, 2008). Organic carbon content was determined using the loss-on-ignition method. 30 g of the dry soil sample was weighedburnt at 380°C for 6 hours, and the fraction of organic carbon content was calculated as previously described (Hoogsteen et al., 2015; Scrimgeour, 2008). Total nitrogen was measured in 50 mg of soil using the FlashSmart CHNS/O elemental analyser (ThermoFischer, Waltham, MA, USA). The standards: BBOT (2,5-Bis (5-tert-butyl-benzoxazol-2-yl) thiophene), Tocopherol Nicotinate and a soil reference material were used to calibrate the instrument.

### 2.3. Chlorophyll concentration and water content

The chlorophyll of each sample was extracted using a protocol based on Castle et al., (2010) and Ritchie, (2006). The extraction was done using methanol, with a soil:methanol ratio of 3:9, followed by a 15-minutes incubation at 65°C and a 2-hour incubation at 4°C. The samples were measured by spectrophotometry at 665 nm and the concentration of chlorophyll was calculated following Ritchie, (2006). Positive controls were spirulina with a concentration of 0.003g/g of soil and negative controls were distilled water. The concentrations are presented in mg of chlorophyll per gram of soil (mg chla/g soil).

### 2.4. RNA extraction and preparation for sequencing

RNA was extracted from 0.5 – 1 g of soil using phenol-chloroform, following the protocol from Angel (2012). The extracted total nucleic acid was treated with DNase to remove the DNA. The remaining RNA was cleaned using parts of the *MagListo* RNA Extraction kit (Bioneer, Daejeon, South Korea). The RNA was reverse transcribed to cDNA using Superscript IV (ThermoFischer, Waltham, MA, USA), and purified using the PCR purification kit (Bioneer, Daejeon, South Korea) in accordance with the manufacturers’ instructions. The cDNA was used as a template to amplify the V3F(341) and V4R(806) regions of the 16S rRNA with CS1/CS2 extensions (Table A.1), in triplicates. Library preparations and sequencing were performed at the Research Resource Centre at the University of Illinois with pair end (2 x 300 bp) MiSeq platform (Illumina, San Diego, CA, USA). Due to low concentrations of ribosomes in the dry soil collected during the summer of 2017, we had to re-extract and re-sequence these samples. However, COVID-19 restrictions prohibit us from using the same sequencing platform, and we were forced to use the facilities and resources available to us at the time. Therefore, RNA was extracted using the RNeasy PowerSoil Total RNA Kit (Qiagen, Hilden, Germany), following the manufacturer’s protocol. Then, the V3-V4 region of the 16S rRNA was amplified in triplicates using the CS1-V3F(341) and CS2-V4R(515) primers (Table A.1). The samples were sequenced (2 x 150 bp) on the iSeq platform (Illumina, San Diego, CA, USA) at the Central and Northern Arava R&D Center (Israel).

### 2.5. Community analysis

Reads were merged, quality checked, and trimmed following the NeatSeq-Flow pipeline (Sklarz et al., 2018). The sequences were analysed using QIIME2 (Bolyen et al., 2018) and Dada2 (Callahan et al., 2016). Reads were clustered in amplicon sequence variants (ASVs) and taxonomy was assigned using Silva v138 (Quast et al., 2013). The total number of sequences can be found in Table A.2. All raw sequences used in this study can be found in BioProject (https://www.ncbi.nlm.nih.gov/ bioproject) under the submission number PRJNA718159.

### 2.6. Functional predictions

Functional predictions of the 16S amplicons was done using Piphillin (Iwai et al., 2016; Narayan et al., 2020) and the KEGG database with a 97%-identity cut-off (May 2020) (Kaneshisa and Goto, 2000). Steps of metabolic pathways for different methods of harvesting energy (organotrophy, lithotrophy and phototrophy) (Cordero et al., 2019; Greening et al., 2016; León-Sobrino et al., 2019; Tveit et al., 2019), for parts of the nitrogen cycle (Madigan et al., 2009), and for the survival of the individual during a drought (DNA conservation and repair, sporulation and Reactive Oxygen Species (ROS)-damage prevention) (Borisov et al., 2013; Hansen et al., 2007; Henrikus et al., 2018; Preiss, 1984; Preiss and Sivak, 1999; Rajeev et al., 2013; Repar et al., 2012; Slade and Radman, 2011) were selected. Then, we picked out genes of interest from each step in the KEGG database and built our own database (Table A.3). The assignment of function to the KEGG numbers of the abundance table from Piphillin was done in R using phyloseq (McMurdie et al., 2017). The significance of temporal differences in predicted functionalities was evaluated using a non-parametric test (Kruskal-Wallis test and a post-hoc Dunn test (Dinno, 2017; Dunn, 1964; Kruskal and Wallis, 1952).

### 2.7. Statistical analysis

All statistical analysis was done using R (R Core Team, 2016) using the phyloseq (McMurdie et al., 2017) along with the ggplot2 (Wickham, 2016), vegan (Oksanen et al., 2014), magritt (Wickham and Bache, 2014), dplyr (Wickham et al., 2018), scales (Wickham, 2017), grid (Murrell, 2004) packages. The significance of difference between time points was determined using a non-parametric test: Kruskal-Wallis test and Dunn test (Dinno, 2017; Dunn, 1964; Kruskal and Wallis, 1952).

## 3. RESULTS

### 3.1. Temporal changes in the biocrust chlorophyll and chemical analyses

We have followed changes in the biocrust before, during and after a rain event and noted that a day after the rain (at T[1]) the biocrust in the sampling site was visibly greener than at any other sampling point (Figure 1). Following our observations, Figure 2 depicts the average chlorophyll concentrations along with the soil water content in the biocrust at each sampling point (Table A.4). The biocrust water content significantly increased between the dry season T[0] and the rain event T[R] (2.26% and 16.2%, respectively, *p* = 0.05; Table A.5). Then soil moisture significantly decreased to 3.67% at T[3] (*p* < 0.05). The chlorophyll concentrations significantly increased right after the rain event (from 8.45 mg chl/g soil to 14.57 mg chl/g soil, during the rain event, *p* = 0.0002; Table A.5), but decreased significantly in later days (from 14.57 mg chl/g to 11.17 mg chl/g soil, three days after the rain, *p* > 0.02; Table A.5). The total organic carbon (Figure A.1) and total nitrogen (Figure A.2) showed slight temporal changes (Table A.4) that were not significant (Table A.5).

### 3.2. Temporal changes in the microbial community composition

Figure 3 shows the bacterial community composition at the order level for each sampling point. The community is mostly composed of *Cyanobacteria, Actinobacteria*, and *Proteobacteria* (Figure 3; Table A.6). During the dry season, biocrust community composition differed significantly from the community depicted during the rain event (Table A.7). The differences were shown mostly in orders belonging to the *Actinobacteria* and *Cyanobacteria* phyla (Figure 3; *p* < 0.05, Table A.7). The relative abundance of *Cyanobacteria*, dominated by the *Cyanobacteriales*, increased during the rain event (from 22% to 41%, Table A.6; *p* < 0.05, Table A.7). While the relative abundance of *Actinobacteria*, dominated by *Micrococcales*, decreased during the rain event (from 50% to 19%, Table A.6; *p* < 0.05, Table A.7). In the days following the rain event, no major changes were detected in the biocrust community (Figure 3; Table A.6 and A.7).

### 3.3. Temporal changes in the microbial function

Figure 4 shows the predicted function based on the taxonomic composition using Piphillin (Iwai et al., 2016; Narayan et al., 2020). The values of the putative functional genes are displayed in copy number (CN). The values were mostly lower in the dry season compared to the hydration-desiccation cycle, except for light and energy sensing (Figure 4; Table A.8). These differences were shown to be statistically significant (*p* < 0.03; Table A.9).

## 4. DISCUSSION

In the Negev Desert, biocrust bacterial communities were shown to alter during hydration. Most apparent was the change in the relative abundance of *Cyanobacteria* which increased while the abundance of *Actinobacteria* decreased (Figure 3) similar to results obtained under controlled conditions that hydrated the biocrust to saturation (Angel and Conrad, 2013). In another controlled experiment, the filamentous cyanobacterium *Leptolyngbya sp.* isolated from desert biocrust, was shown to respond quickly to both hydration and desiccation (Oren et al., 2019, 2017). Slight increase in biocrust moisture, triggered by dew simulation, induced DNA repair and associated regulatory genes for activating the photosynthetic system of the cyanobacterium (Murik et al., 2017; Rajeev et al., 2013). However, in this study we show that during a rain event of the Negev Desert biocrust, soil moisture increased and resulted in: (i) a sharp rise in chlorophyll concentrations (Figure 2), (ii) a significant increase in the relative abundance of various cyanobacterial orders (Figure 3; Table A.6 and A.7), and (iii) a significant increase in putative phototrophy of the community (Figure 4; Table A.8 and A.9). The concentration of the chlorophyll pigment was suggested to be linked to the water content (Péli et al., 2011) and to the activity of dominating primary producer in the biocrust, i.e., *Cyanobacteria* and/or green algae (Madigan et al., 2009). While cyanobacterial activity increased with soil moisture, no significant changes were detected in the total organic carbon and nitrogen content (Figure A.1 and A.2; Table A.4 and S5). This observation suggests that the immediate change in these parameters is negligible compared to existing soil reservoir; thus, it cannot be used as an indicator for the reaction of the local community to rain events. Moreover, it was recently proposed that in arid biocrusts, the dominant *Cyanobacteria* exchange carbon for nitrogen with copiotrophic diazotrophs, thus rapidly utilizing available nutrients and enabling the colonisation of the oligotrophic dryland soils (Couradeau et al., 2019).

In desert soil, rain events were shown to entail a decrease in the abundance of *Actinobacteria* both in the biocrust (Angel and Conrad, 2013) and topsoil (Št’ovíček et al., 2017). Members of this phylum were shown to be well adapted to living in harsh environments (Goodfellow and Williams, 1983; Zvyagintsev et al., 2007), and were found to be abundant in the Negev Highland biocrust (Meier et al., 2021). Here we showed that the increase of water content may lead to an increase in activity in all gene groups linked to energy usage or production (Figure 4; Table A.8). The generally dry biocrust experiences a narrow window of hydration conditions after a rain event of only a couple of days (Figure 2). These conditions are rapidly exploited by phototrophs before the soil dries (Figure 2 and 3), and inhibit resilient heterotrophs (Figure 3), as was previously shown in controlled (Cordero et al., 2019; Greening et al., 2016; León-Sobrino et al., 2019; Tveit et al., 2019), and naturel (León-Sobrino et al., 2019) settings.

We showed that re-hydration stage is quick and bacterial activity restarts within hours. However, during the desiccation stage, changes are slower. The biocrust dries quickly after the rain (Figure 2) due to evaporation, encouraged by the strong radiation, high winds, and low air humidity (Kidron and Tal, 2012). Yet, unlike the response to dew hydration (Oren et al., 2019, 2017), the community does not immediately inactivates. We sampled the soil along the hydration-desiccation cycle and stopped sampling when the soil was dehydrated because we expected inactivation of the community. In a previous study (Št’ovíček et al., 2017), we showed that the topsoil community bounces back to its original structure as soon as the soil dries. Yet in the biocrust, dehydration was associated by a decrease in chlorophyll concentrations, yet we detected no significant changes in the community composition (Figure 3). This may be possible due to the EPS produced by the *Cyanobacteria* (Mager and Thomas, 2011; Mazor et al., 1996) that dominate the biocrust. *Cyanobacteria* were shown to secrete copious amounts of EPS that bind the soil particles together (Kidron et al., 2020; Kidron and Tal, 2012). The EPS in the biocrust was shown to retain water in the soil, slowing down the drying process (Roberson and Firestone, 1992). Likewise, EPS in soil was shown to create microhabitats that retain humidity (Colica et al., 2014), thus stabilising the biocrust (Lan et al., 2012), and protecting the residing microorganisms from desiccation (Mazor et al., 1996). EPS is a key component in the Negev Desert biocrusts (Kidron et al., 2020). Here we propose that EPS may benefit the microbial community by creating microhabitats in which moisture is retained longer, enabling extended active phase following a rain event. This extra time to propagate after a rain event, may justify the ample resources invested by the *Cyanobacteria* in EPS production (Mager and Thomas, 2011). To validate this hypothesis further study is required.

## 5. CONCLUSION

In desert biocrusts, bacterial communities must respond quickly to hydration, in order to take advantage of the short windows of opportunity to photosynthesize, and then to desiccation to prevent cells damage. Our findings reinforce controlled studies showing that biocrust hydration change the bacterial community, increasing the *Cyanobacteria* abundance over *Actinobacteria*. However, here we have shown that under field conditions the response to soil desiccation is slower, allowing for a longer period of activity and production even a day after soil moisture decrease. we speculate that the lag in response to dehydration is due to EPS-based water retention in the soil mediated by the *Cyanobacteria* producers, justifying the metabolic cost of biocrust formation.

## ACKNOWLEDGEMENT

The authors are grateful to Lusine Ghazaryan for technical support and to Ben Poodiack for editing the manuscript. This study was partially supported by the Israel Science Academy, grant no. 993/11.

## APPPENDIX A

**Table A.1.**
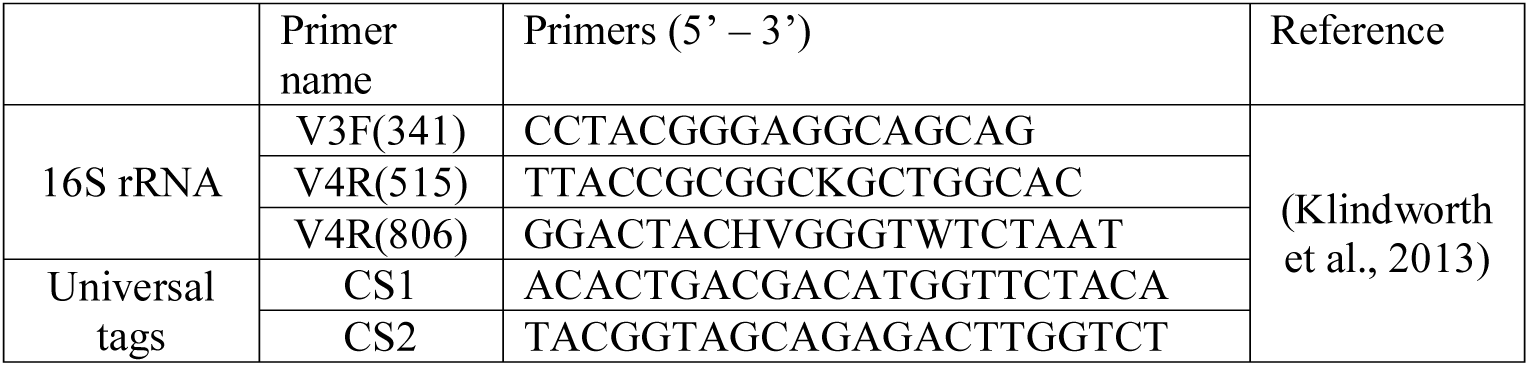
Primers used in this study.

**Table A.2.** Statistics from Dada2.

| Sample<br>(Time<br>point) | Input | Filtered | Percentage<br>of input<br>passed<br>filter | Denoised | Non-<br>chimeric | Percentage<br>of input<br>non-<br>chimeric |
| --- | --- | --- | --- | --- | --- | --- |
| T[R] | 99090 | 87403 | 88.21 | 76272 | 68762 | 69.39 |
| T[R] | 102014 | 87207 | 85.49 | 75796 | 64954 | 63.67 |
| T[R] | 107763 | 94407 | 87.61 | 80242 | 72676 | 67.44 |
| T[R] | 94175 | 81352 | 86.38 | 69460 | 61519 | 65.32 |
| T[R] | 97752 | 85658 | 87.63 | 76694 | 65590 | 67.10 |
| T[1] | 102147 | 89670 | 87.79 | 79436 | 68611 | 67.17 |
| T[1] | 110406 | 96638 | 87.53 | 86745 | 76384 | 69.18 |
| T[1] | 94247 | 81576 | 86.56 | 72289 | 65755 | 69.77 |
| T[1] | 107731 | 94180 | 87.42 | 83504 | 72831 | 67.60 |
| T[1] | 96982 | 84993 | 87.64 | 77197 | 67547 | 69.65 |
| T[2] | 95525 | 82453 | 86.32 | 73811 | 63892 | 66.89 |
| T[2] | 90500 | 79303 | 87.63 | 75977 | 74636 | 82.47 |
| T[2] | 84648 | 74376 | 87.87 | 71060 | 69017 | 81.53 |
| T[2] | 96778 | 85143 | 87.98 | 75971 | 66483 | 68.70 |
| T[2] | 83749 | 72395 | 86.44 | 65649 | 60857 | 72.67 |
| T[3] | 85527 | 74977 | 87.66 | 66324 | 56872 | 66.50 |
| T[3] | 92648 | 81056 | 87.49 | 74512 | 67015 | 72.33 |
| T[3] | 98388 | 86526 | 87.94 | 78048 | 69910 | 71.06 |
| T[3] | 92219 | 79938 | 86.68 | 69799 | 62666 | 67.95 |
| T[3] | 88140 | 77515 | 87.95 | 73273 | 72113 | 81.82 |
| T[0] | 22095 | 21646 | 97.97 | 19900 | 12628 | 57.15 |
| T[0] | 23457 | 22888 | 97.57 | 18342 | 11627 | 49.57 |
| T[0] | 26072 | 25368 | 97.30 | 20726 | 12867 | 49.35 |

**Table A.3.**
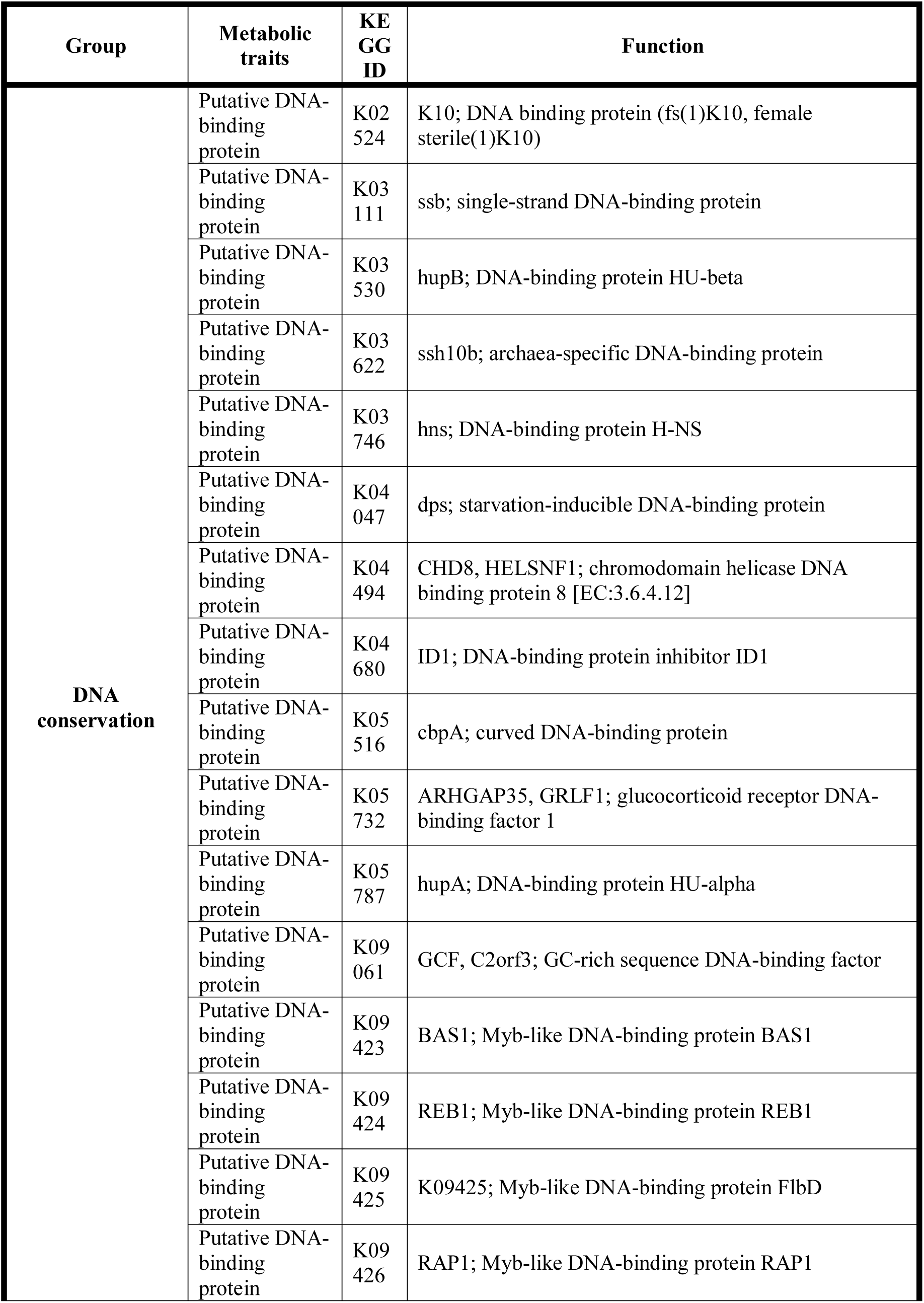
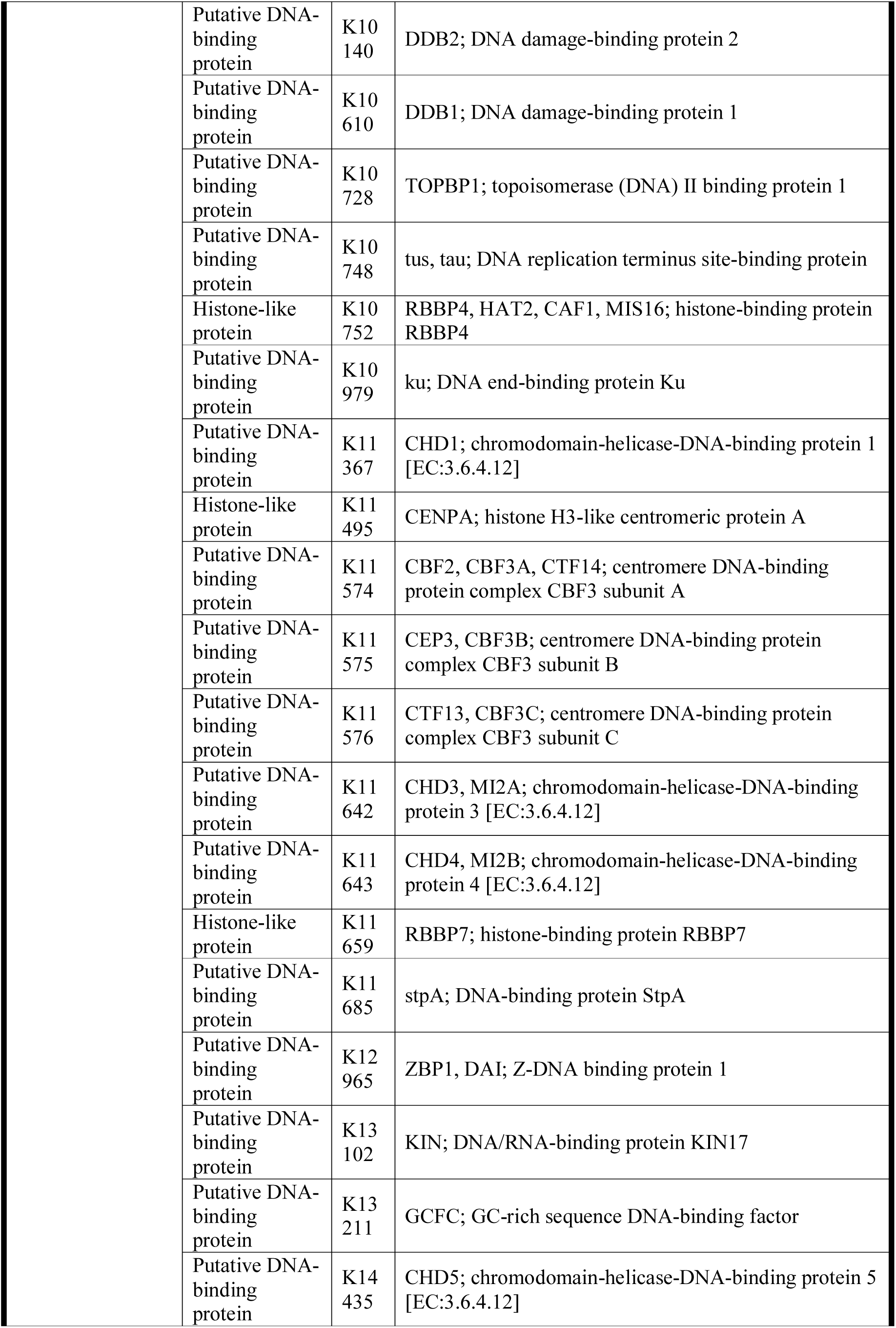
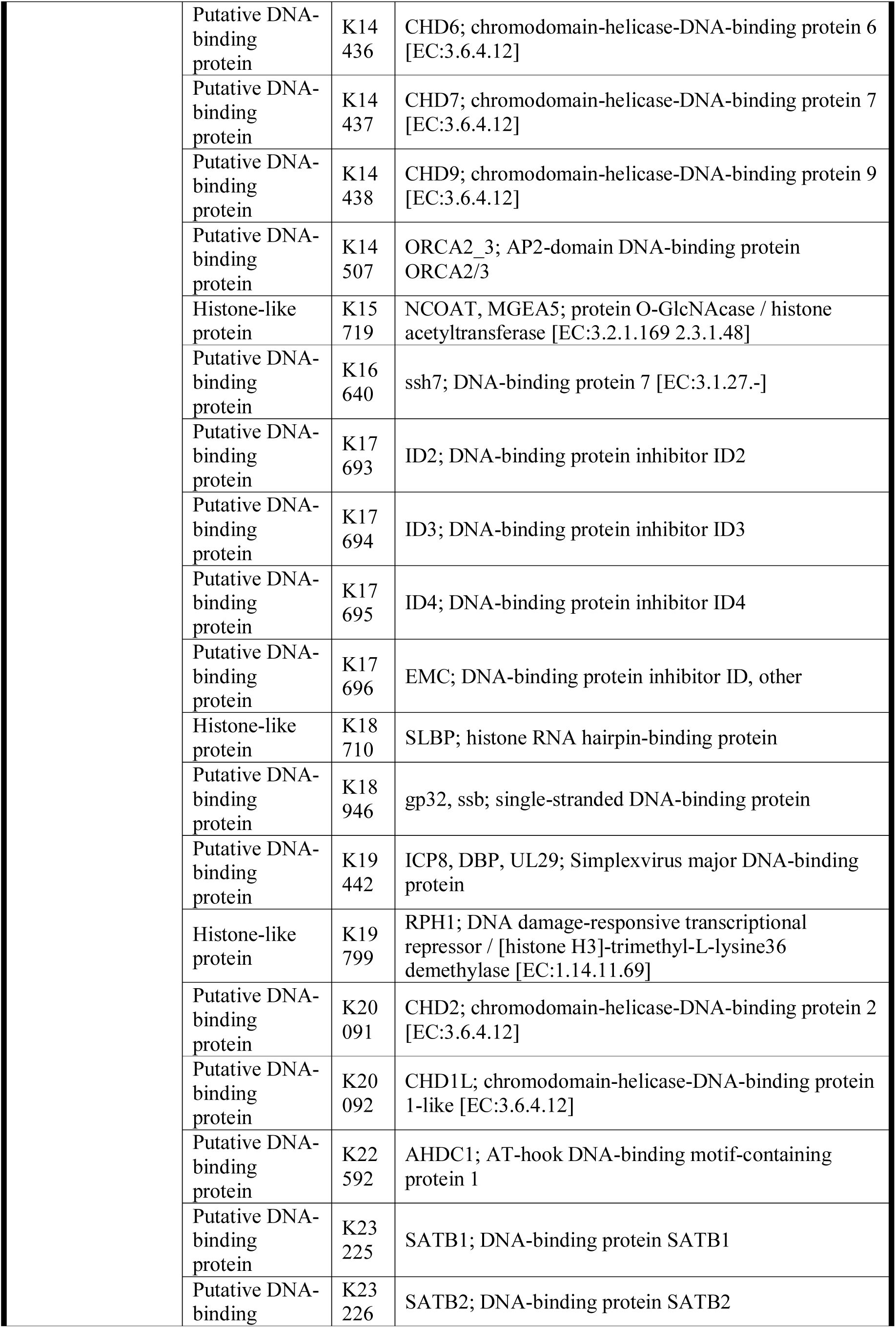
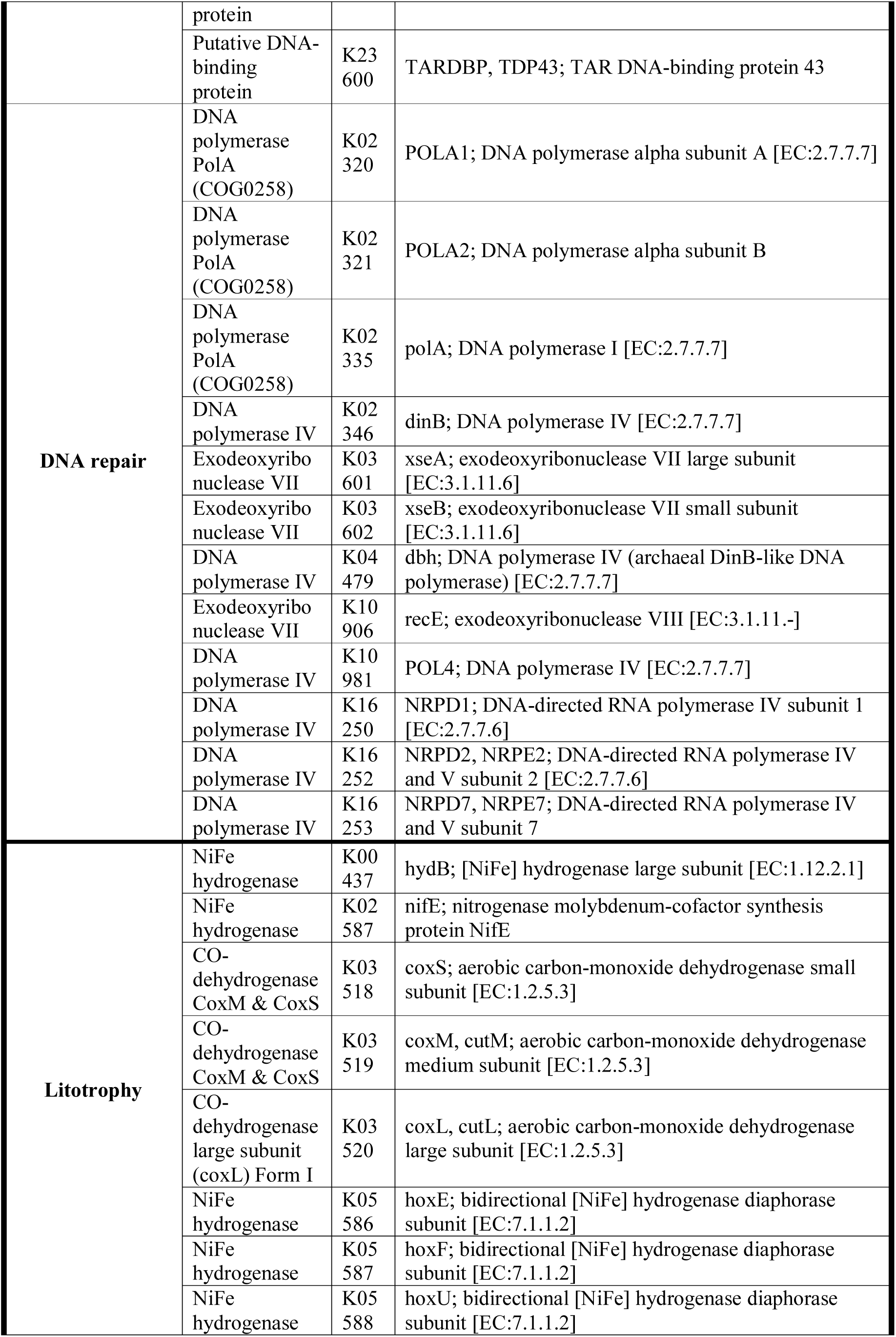
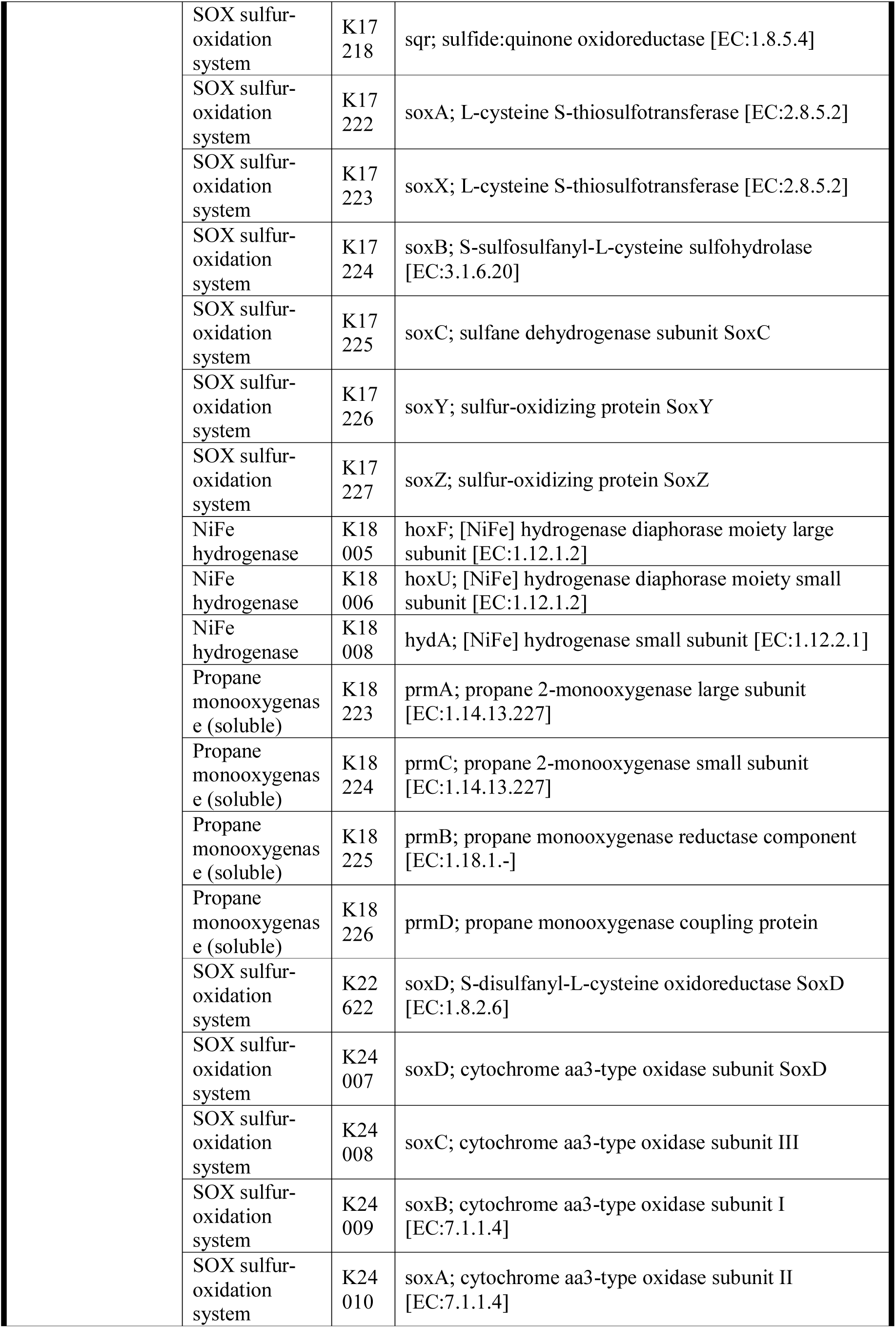
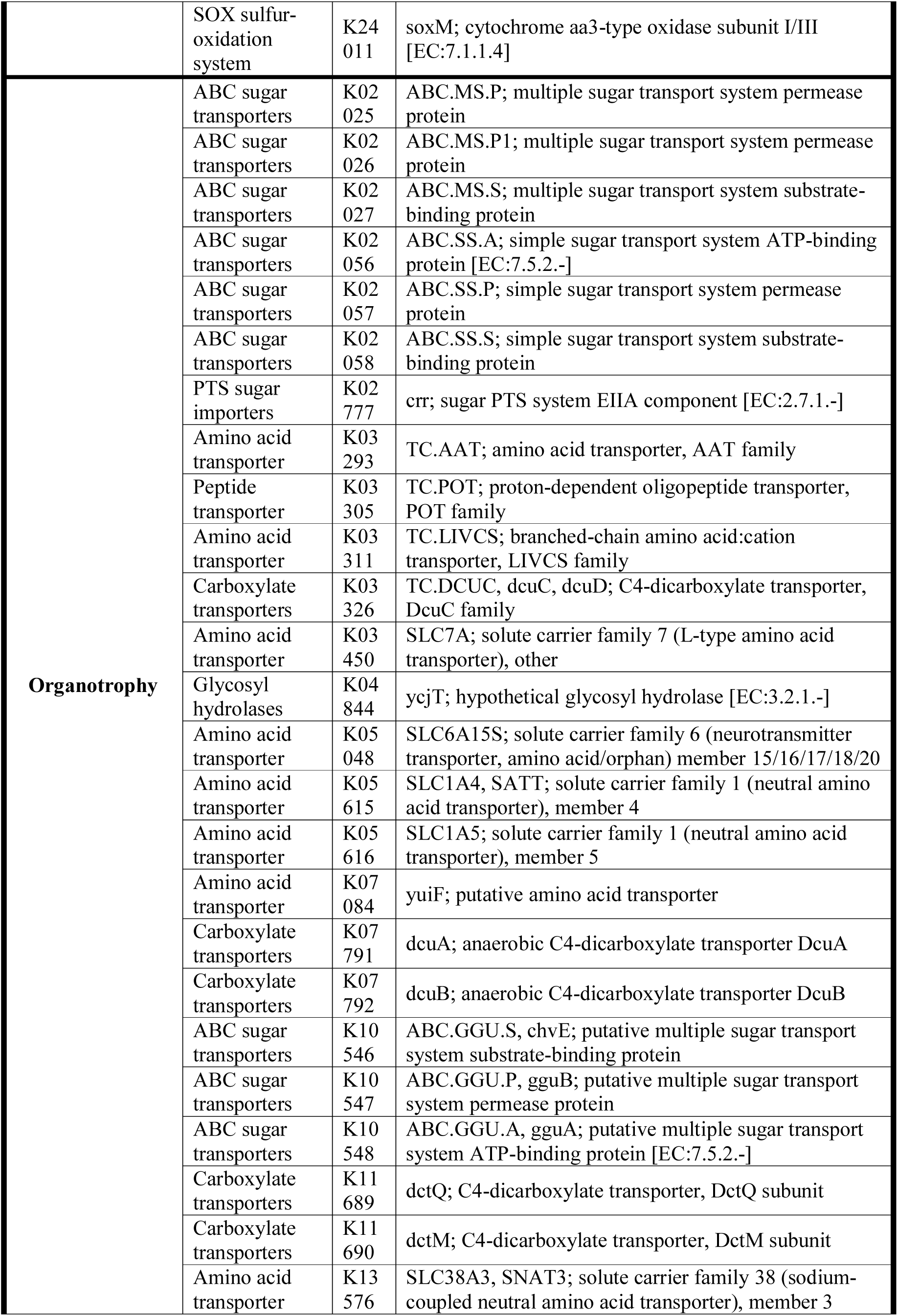
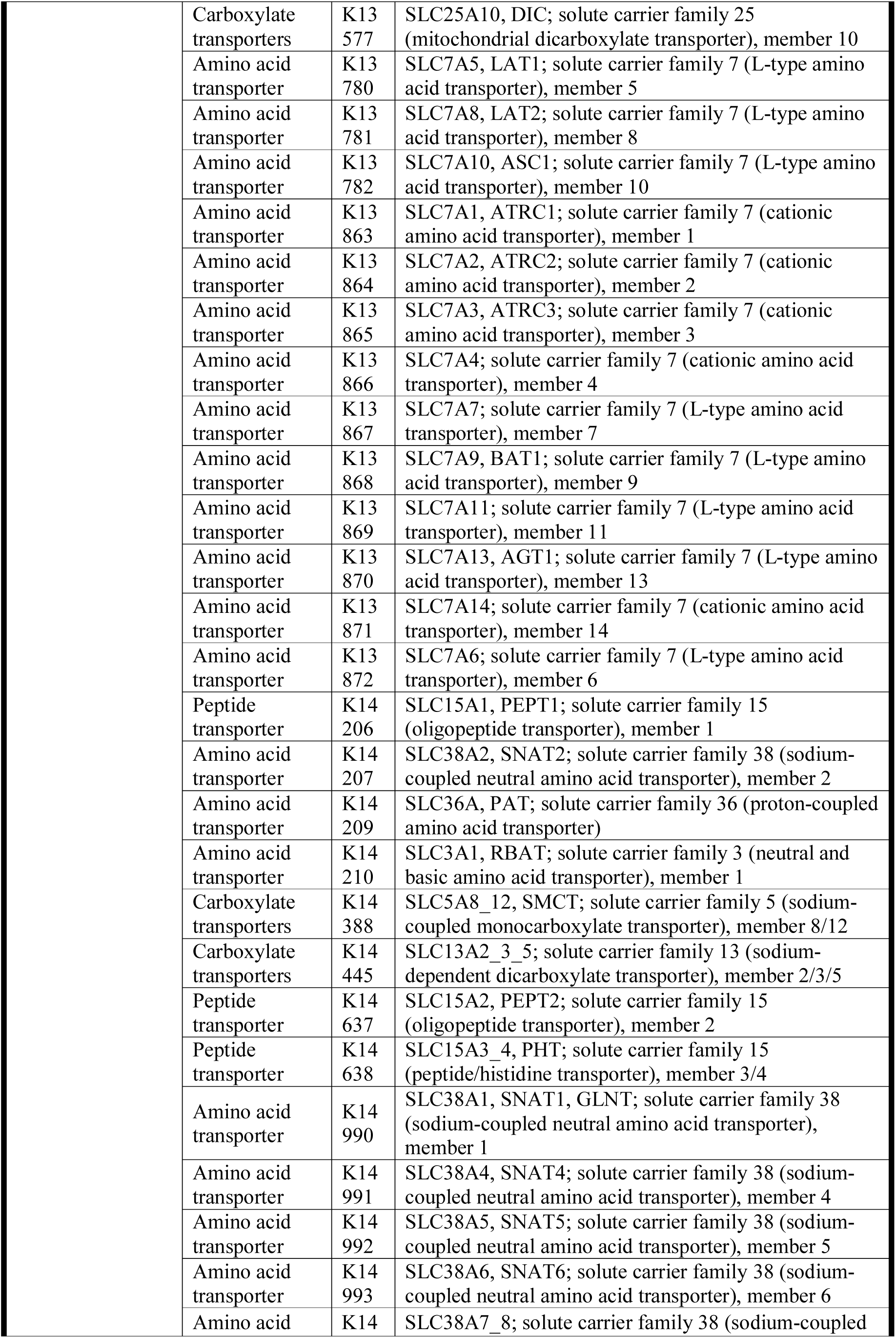
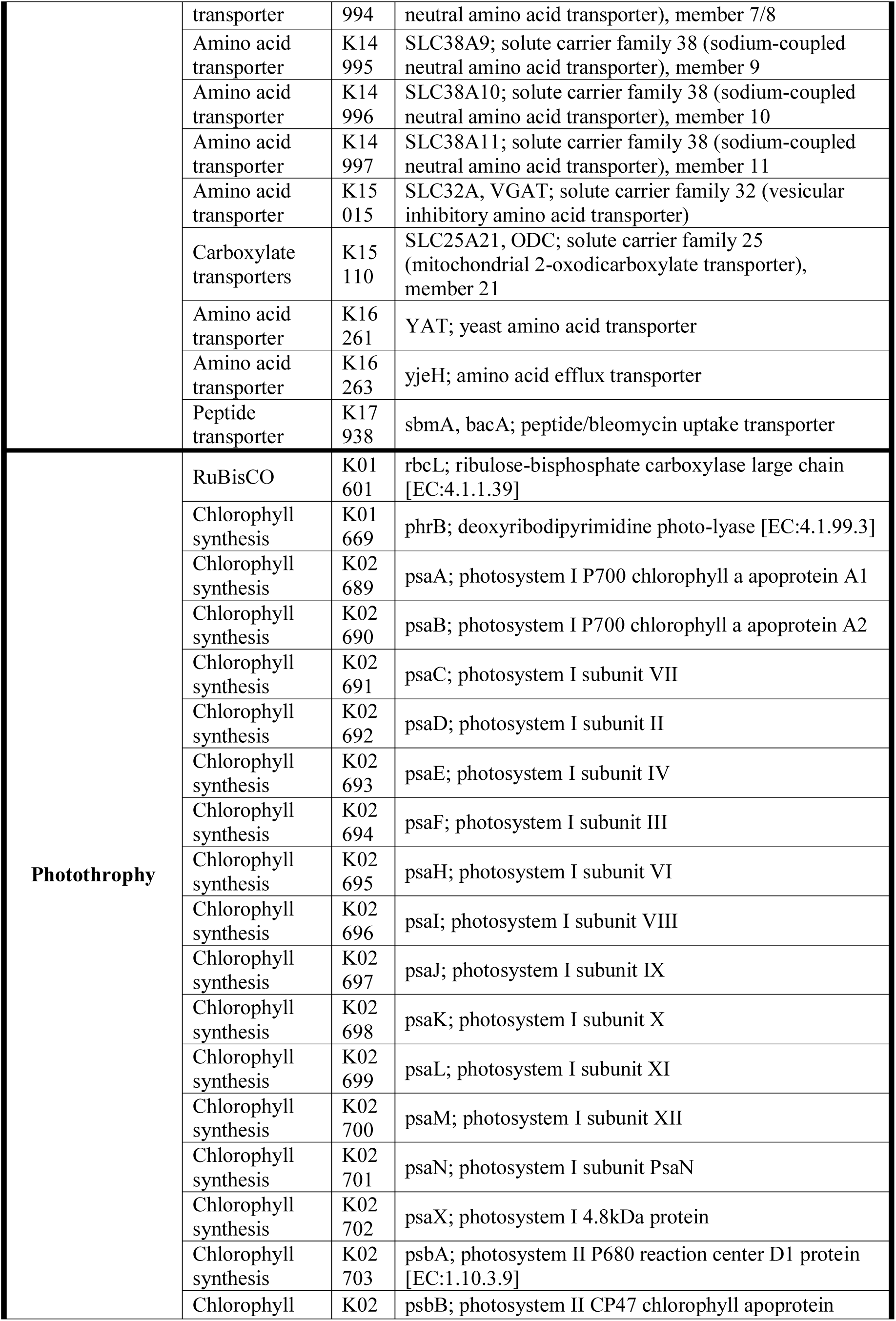
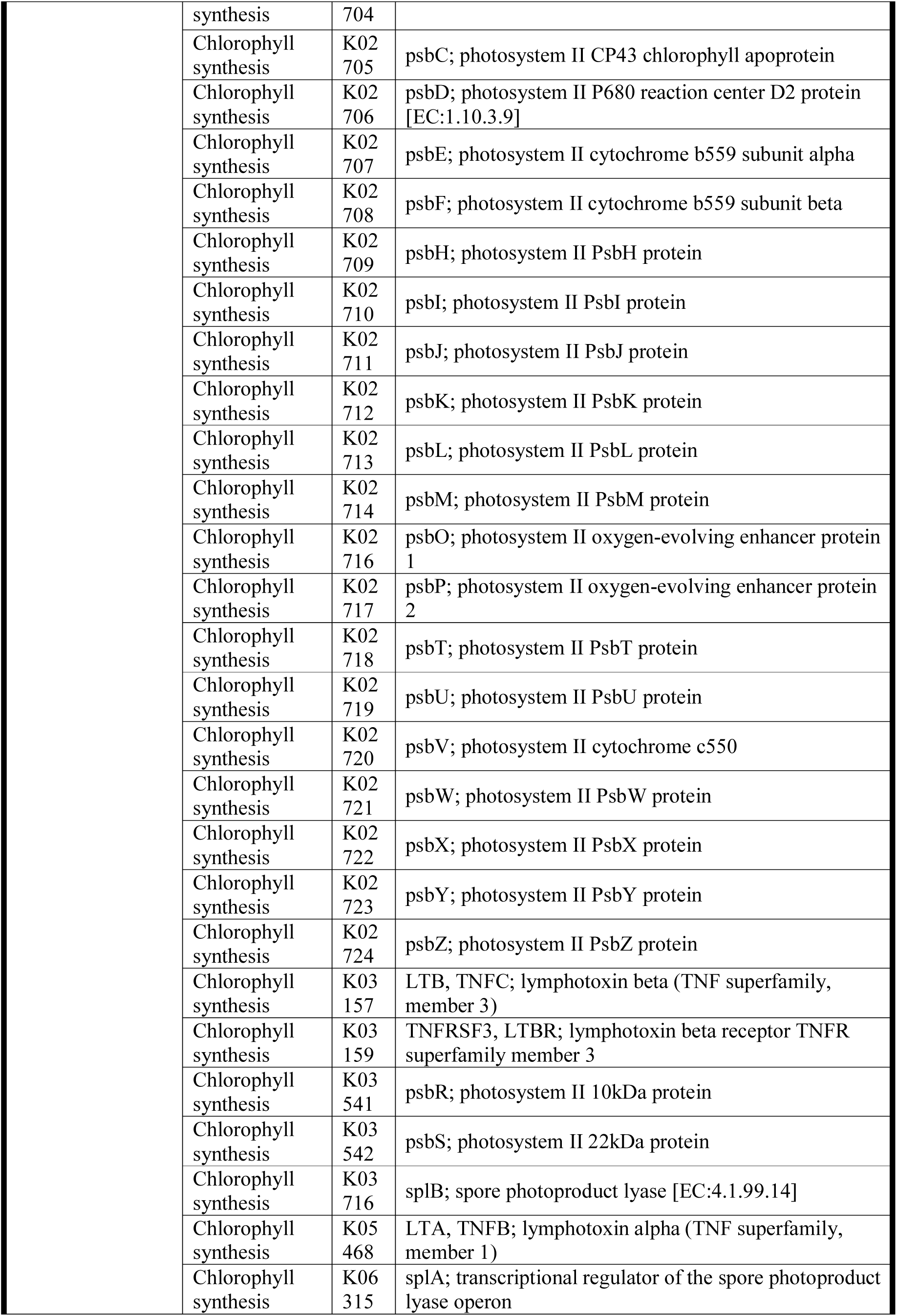
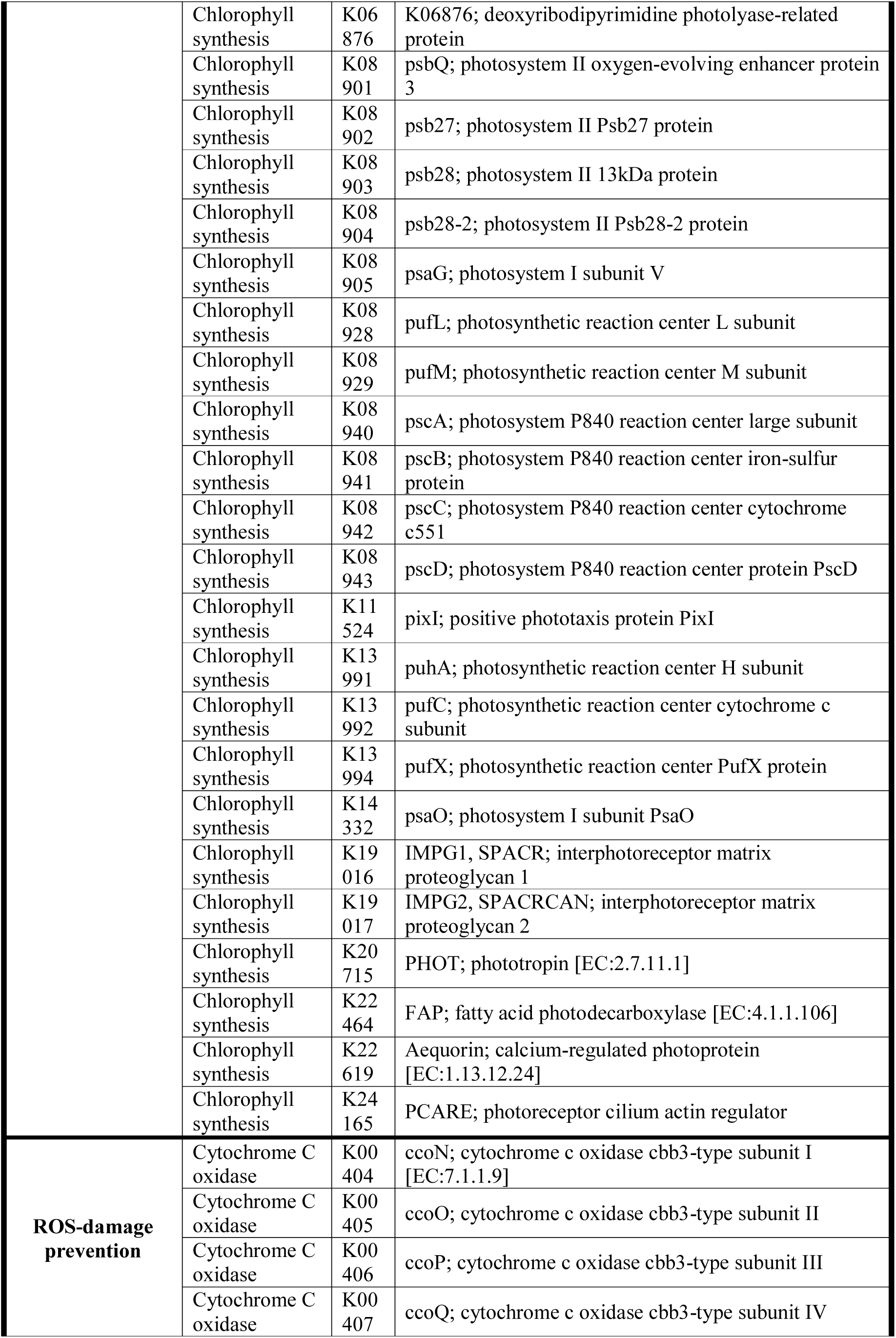
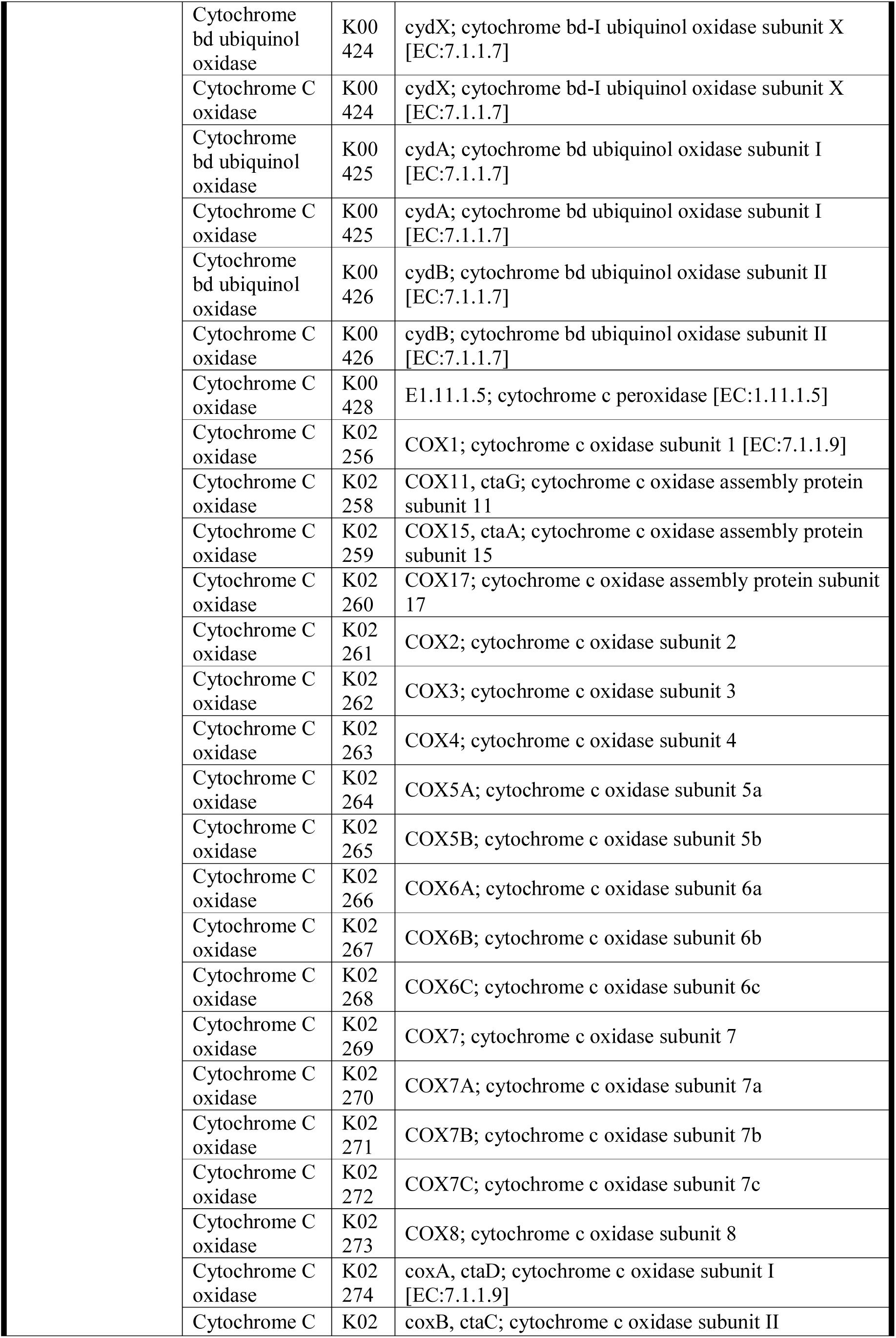
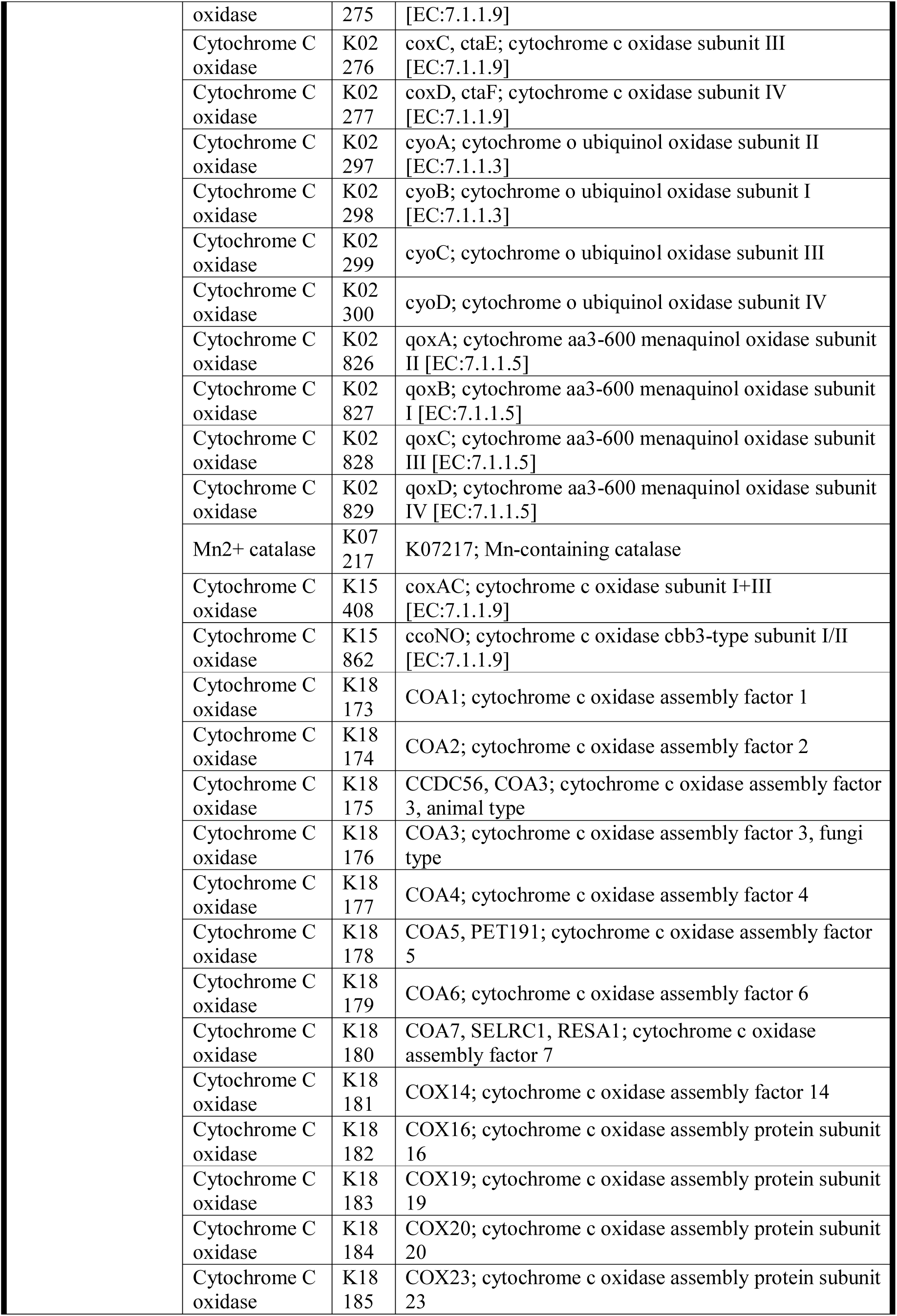
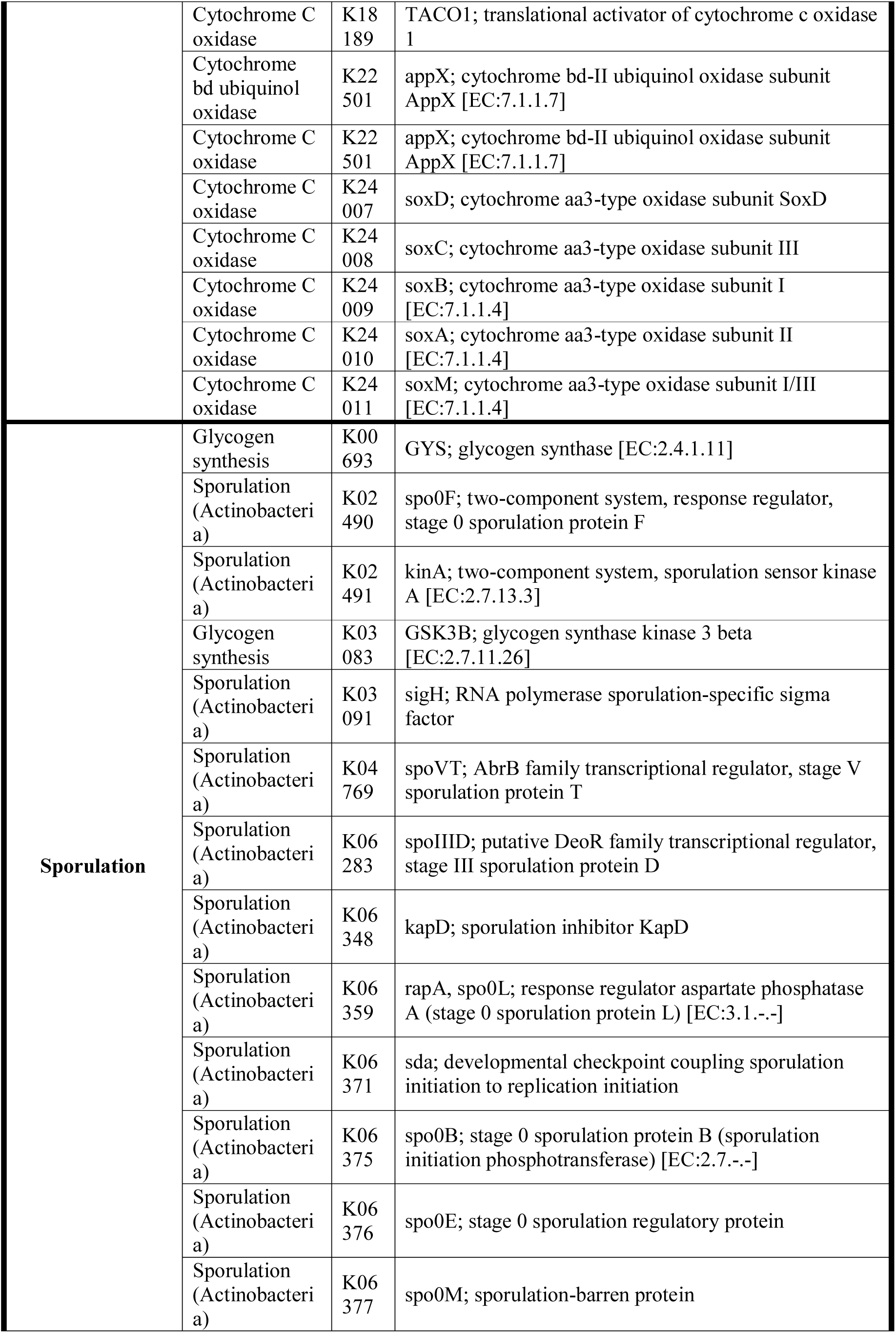
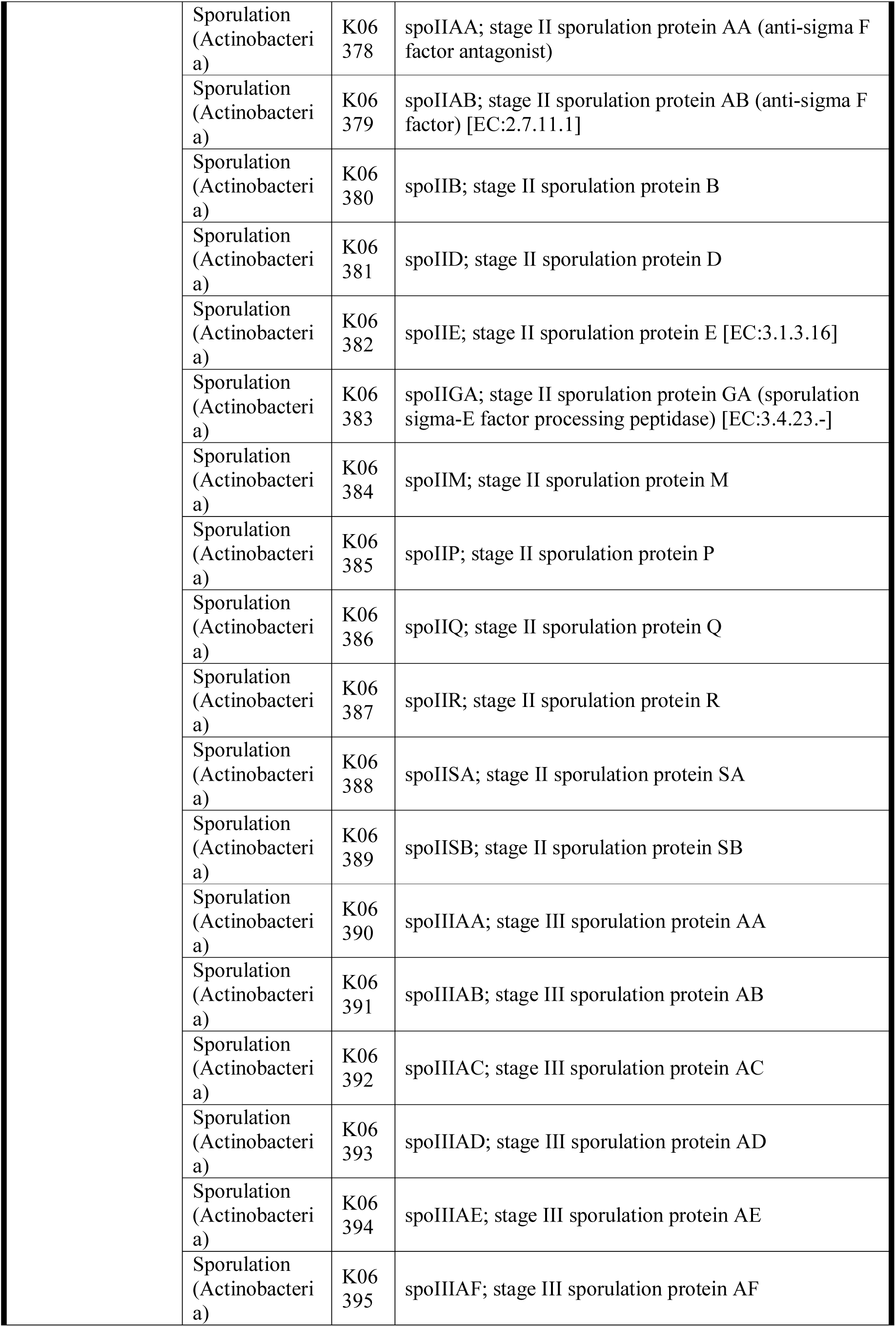
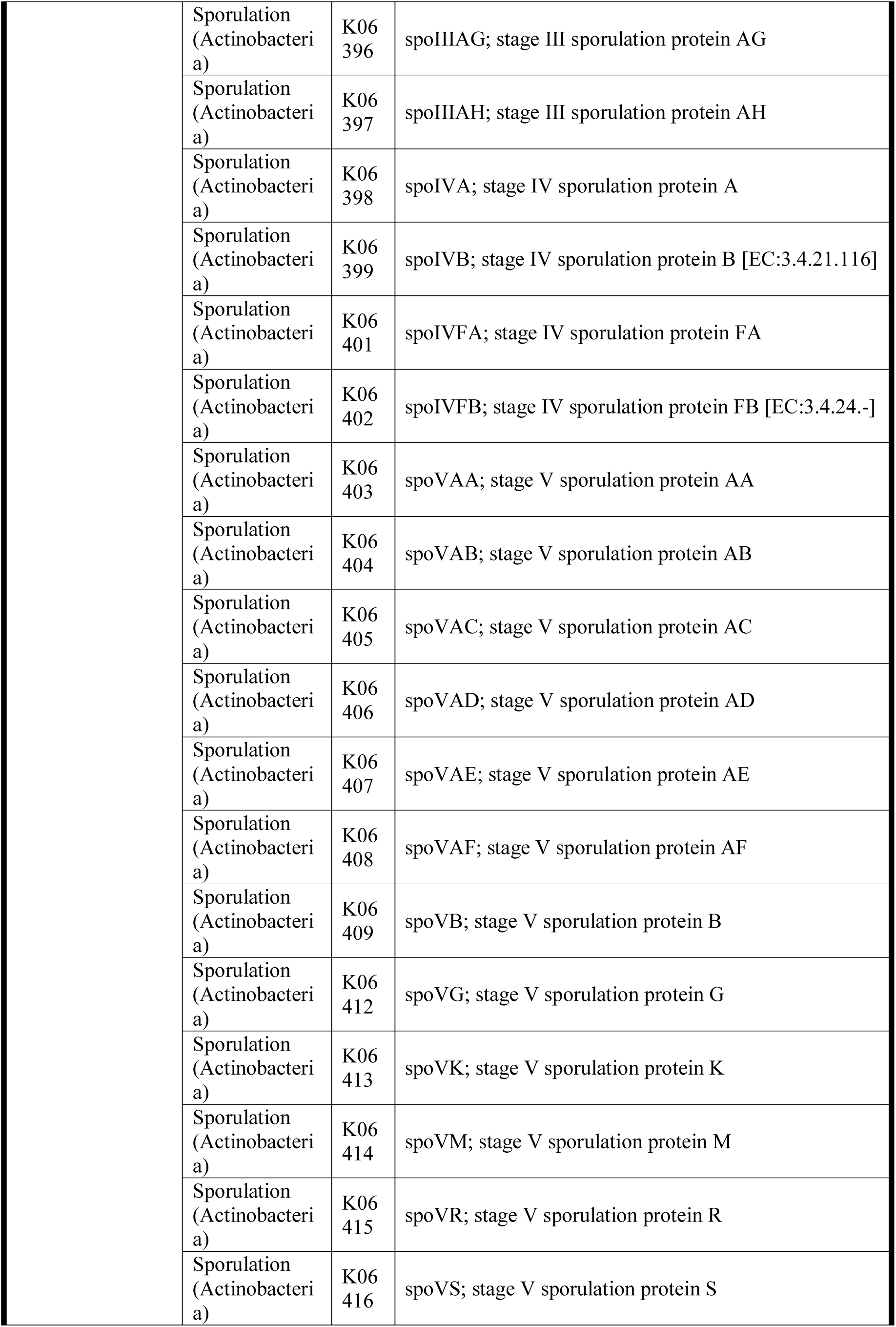
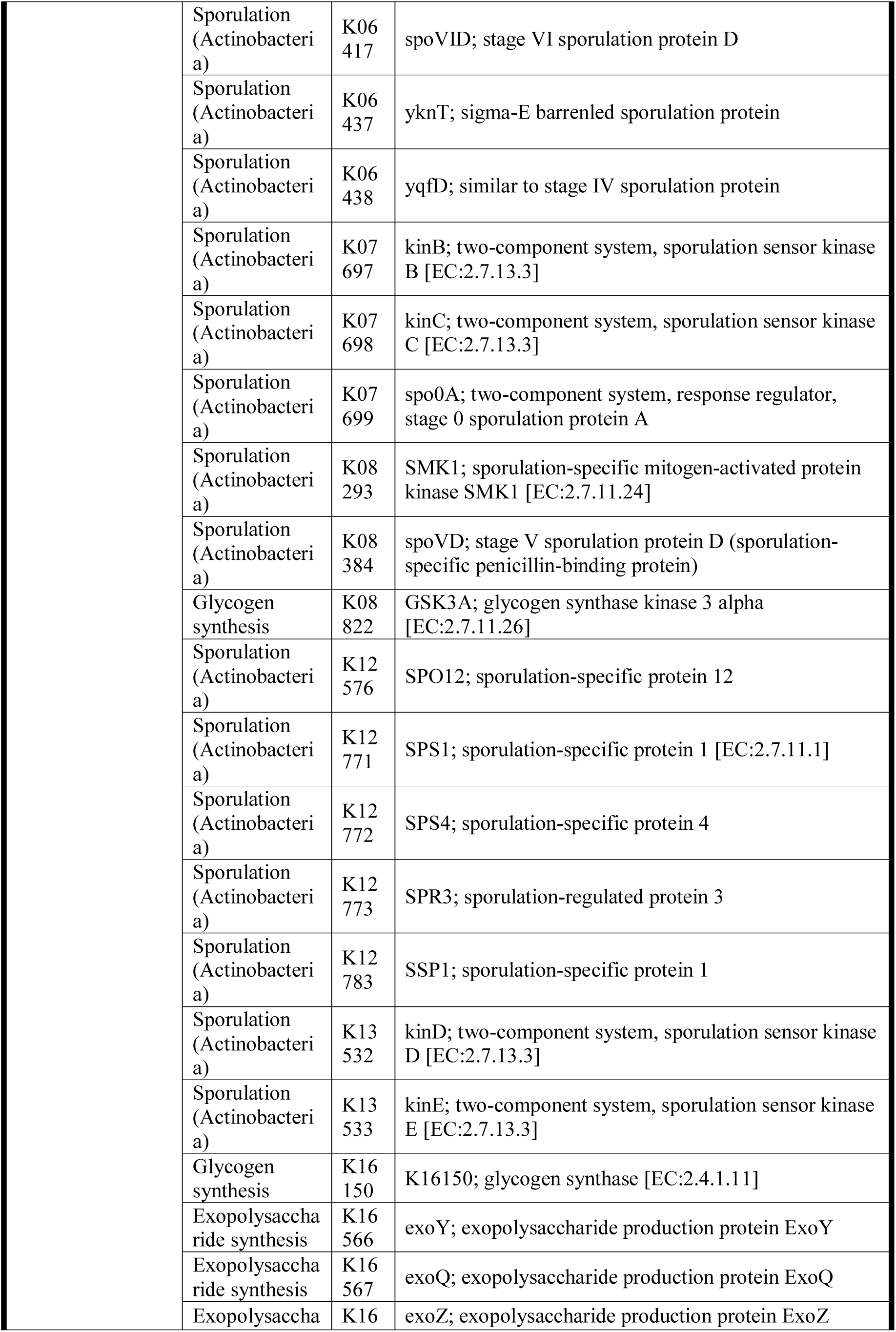
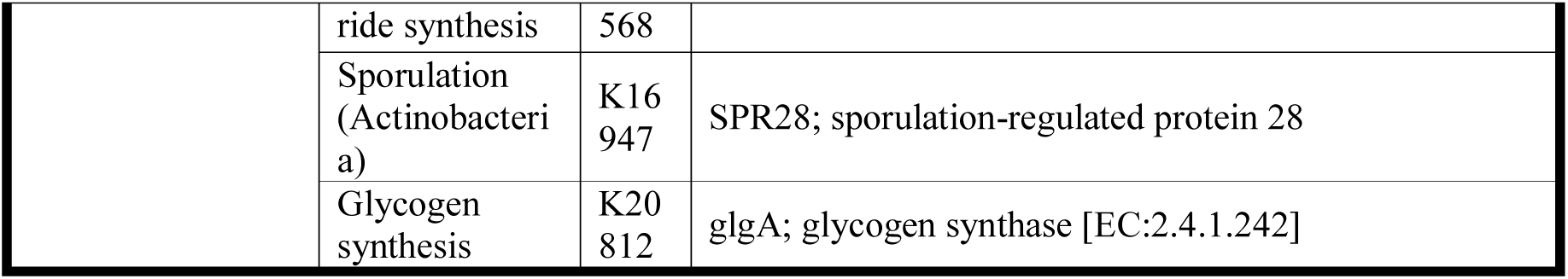
List of the genes used for function prediction ordered by groups and subgroups.

**Table A.4.** Chlorophyll concentrations and water content values in the biocrust at each sampling point and site.

| Time | Chlorophyll<br>concentration<br>(mg chla/ g soil) | Water content<br>(%) | Total organic carbon<br>(%) | Total nitrogen<br>(%) |
| --- | --- | --- | --- | --- |
| T[0] | 7.670 | 1.850 | 4.225 | 0.112 |
| T[0] | 8.553 | 2.670 | 4.107 | 0.114 |
| T[0] | 6.835 | 2.290 | 4.188 | 0.107 |
| T[R] | 10.145 | 13.610 | 2.999 | 0.104 |
| T[R] | 16.144 | 16.930 | 3.898 | 0.104 |
| T[R] | 15.703 | 16.550 | 3.787 | 0.109 |
| T[R] | 12.745 | 16.340 | 3.967 | 0.104 |
| T[R] | 14.160 | 17.520 | 3.972 | 0.071 |
| T[1] | 10.519 | 5.300 | 3.269 | 0.065 |
| T[1] | 16.003 | 5.620 | 3.238 | 0.065 |
| T[1] | 16.877 | 7.140 | 3.955 | 0.085 |
| T[1] | 15.262 | 6.220 | 3.964 | 0.073 |
| T[1] | 14.177 | 6.830 | 3.948 | 0.098 |
| T[2] | 10.092 | 3.920 | 3.879 | 0.087 |
| T[2] | 11.829 | 4.490 | 4.031 | 0.082 |
| T[2] | 11.840 | 4.510 | 4.244 | 0.100 |
| T[2] | 13.045 | 5.070 | 4.134 | 0.115 |
| T[2] | 12.975 | 6.900 | 4.050 | 0.095 |
| T[3] | 9.881 | 2.860 | 3.834 | 0.092 |
| T[3] | 12.751 | 3.670 | 3.709 | 0.100 |
| T[3] | 9.314 | 3.710 | 3.706 | 0.076 |
| T[3] | 9.366 | 4.320 | 4.511 | 0.081 |
| T[3] | 14.520 | 3.780 | 4.589 | 0.093 |

**Table A.5.** Dunn tests p values for chlorophyll concentration (mg chla/g of soil), water content (%), total organic carbon content (%) and total nitrogen (%) in biocrust samples collected at the different sampling point. Bold numbers mark significant differences (<0.05).

| Comparison | Chlorophyll | Water content | Total organic Carbon | Total Nitrogen |
| --- | --- | --- | --- | --- |
| T[0] - T[1] | **0.00538004** | 0.05699315 | 0.89 | 0.169 |
| T[0] - T[2] | **0.02846706** | 0.35971877 | 0.64 | 0.525 |
| T[1] - T[2] | 0.25895220 | **0.02620581** | 0.55 | 0.448 |
| T[0] - T[3] | 0.17516547 | 0.20434975 | 0.75 | 0.065 |
| T[1] - T[3] | 0.05299877 | **0.00804845** | 0.85 | 0.611 |
| T[2] - T[3] | 0.16605599 | 0.32025932 | 0.43 | 0.210 |
| T[0] - T[R] | **3.5589E-06** | 0.15725837 | 0.84 | 0.723 |
| T[1] - T[R] | **0.02620581** | 0.28273460 | 0.95 | 0.300 |
| T[2] - T[R] | **0.00485039** | 0.08612648 | 0.51 | 0.776 |
| T[3] - T[R] | **0.00018814** | **0.03347764** | 0.90 | 0.128 |

**Table A.6.**
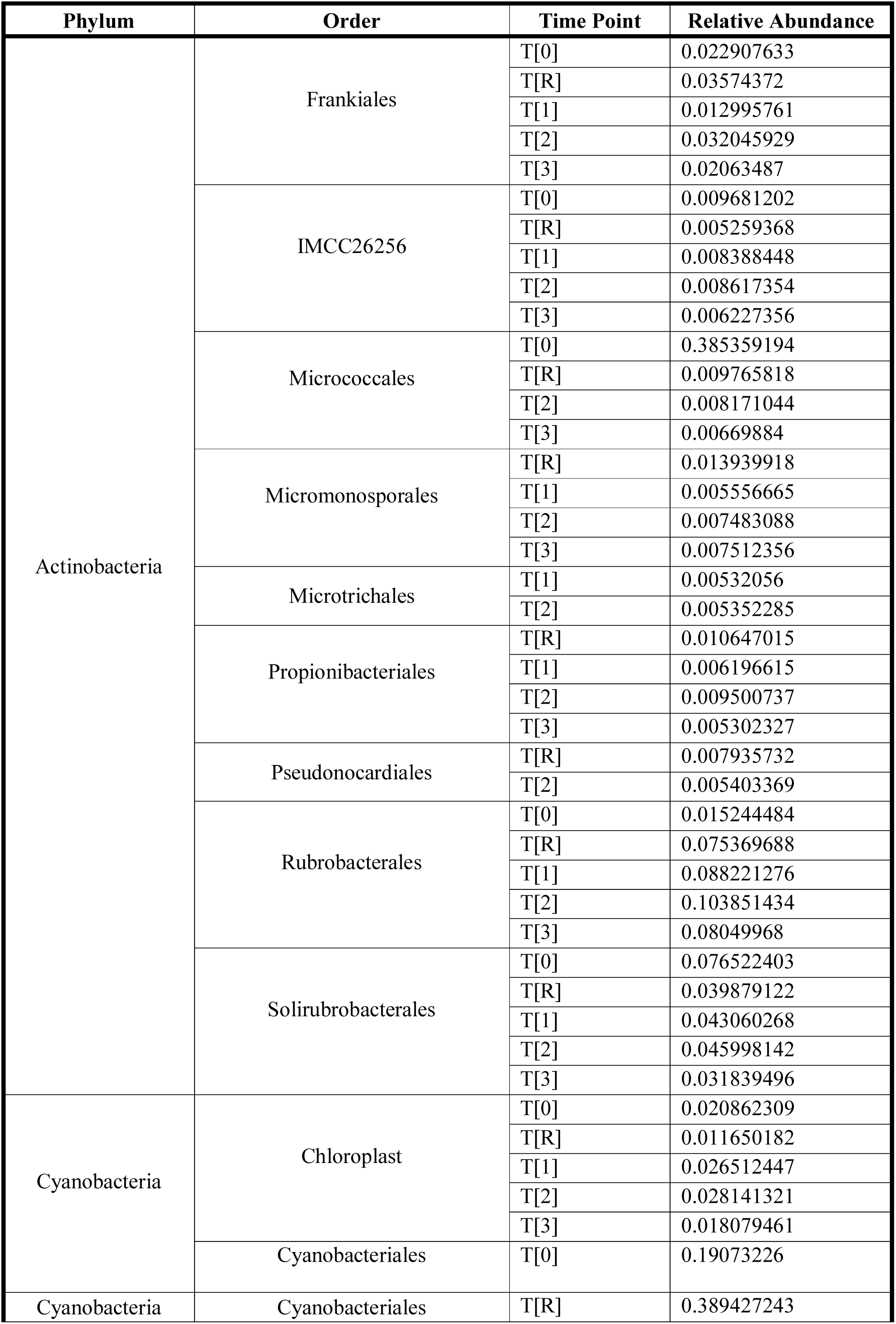
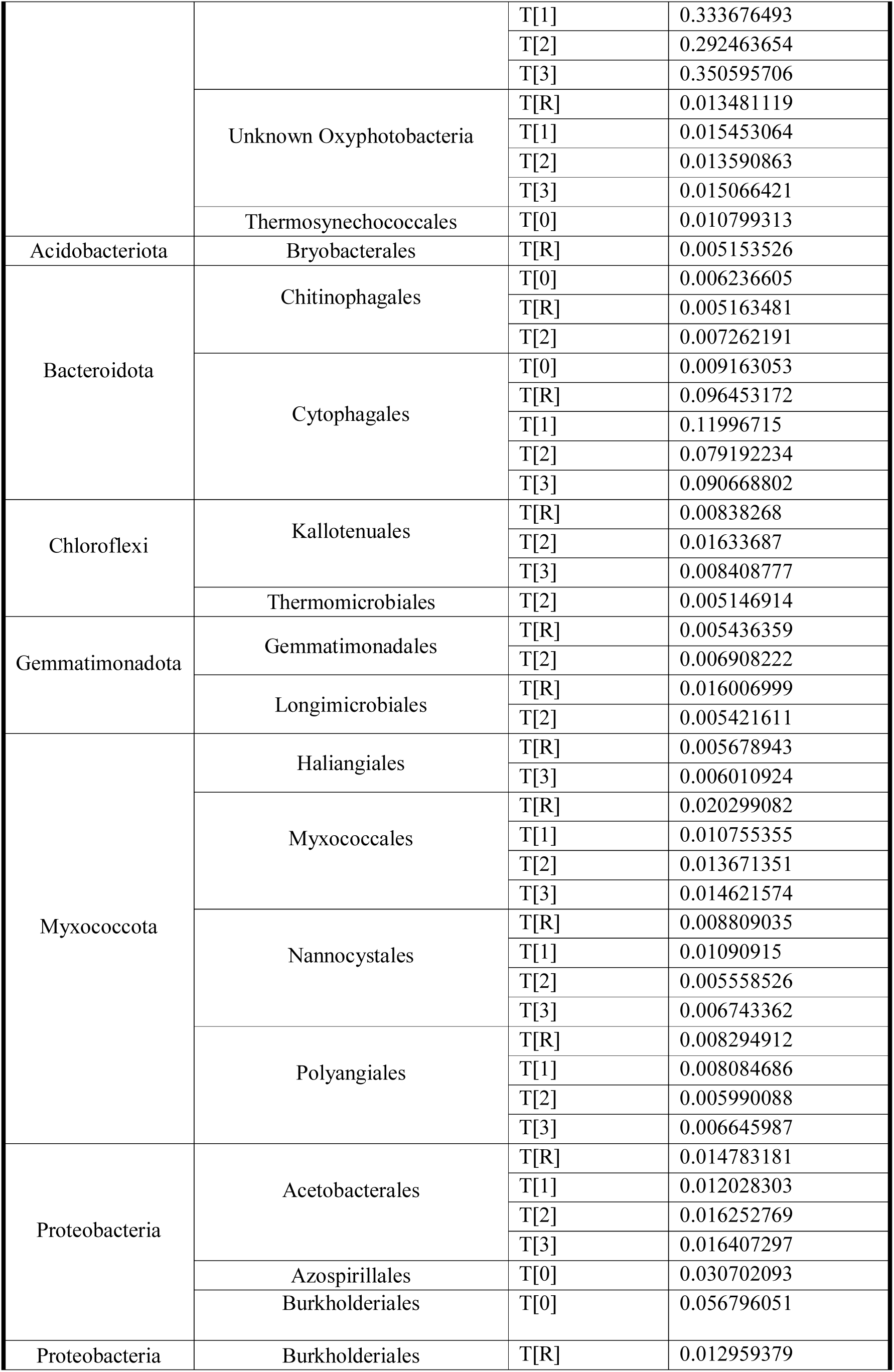
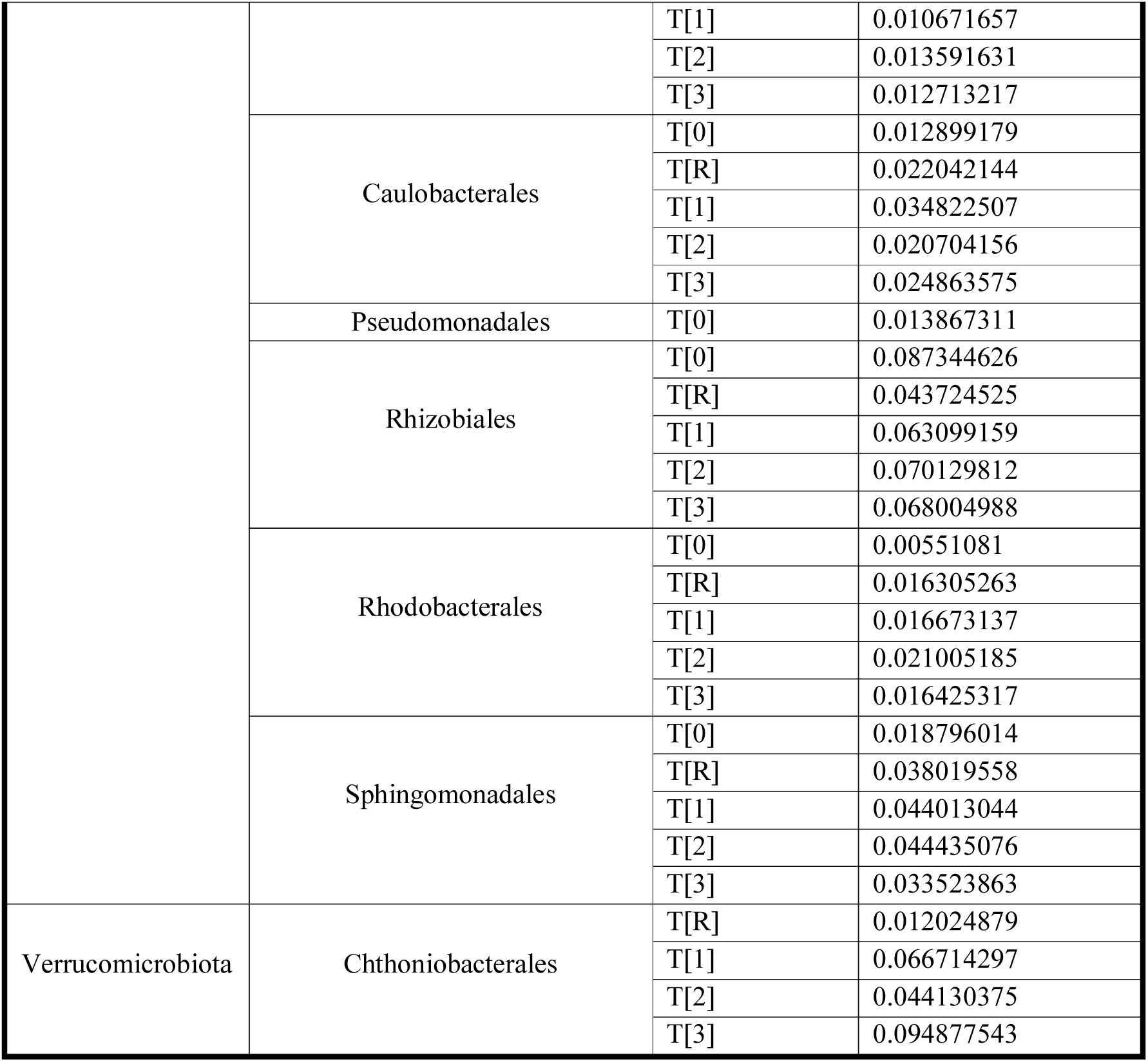
Relative abundance of the taxa in the biocrust community at each time point.

| Phylum | Order | Time Point | Relative Abundance |
| --- | --- | --- | --- |
| Actinobacteria | Frankiales | T[0] | 0.022907633 |
|  |  | T[R] | 0.03574372 |
|  |  | T[1] | 0.012995761 |
|  |  | T[2] | 0.032045929 |
|  |  | T[3] | 0.02063487 |
|  | IMCC26256 | T[0] | 0.009681202 |
|  |  | T[R] | 0.005259368 |
|  |  | T[1] | 0.008388448 |
|  |  | T[2] | 0.008617354 |
|  |  | T[3] | 0.006227356 |
|  | Micrococcales | T[0] | 0.385359194 |
|  |  | T[R] | 0.009765818 |
|  |  | T[2] | 0.008171044 |
|  |  | T[3] | 0.00669884 |
|  | Micromonosporales | T[R] | 0.013939918 |
|  |  | T[1] | 0.005556665 |
|  |  | T[2] | 0.007483088 |
|  |  | T[3] | 0.007512356 |
|  | Microtrichales | T[1] | 0.00532056 |
|  |  | T[2] | 0.005352285 |
|  | Propionibacteriales | T[R] | 0.010647015 |
|  |  | T[1] | 0.006196615 |
|  |  | T[2] | 0.009500737 |
|  |  | T[3] | 0.005302327 |
|  | Pseudonocardiales | T[R] | 0.007935732 |
|  |  | T[2] | 0.005403369 |
|  | Rubrobacterales | T[0] | 0.015244484 |
|  |  | T[R] | 0.075369688 |
|  |  | T[1] | 0.088221276 |
|  |  | T[2] | 0.103851434 |
|  |  | T[3] | 0.08049968 |
|  | Solirubrobacterales | T[0] | 0.076522403 |
|  |  | T[R] | 0.039879122 |
|  |  | T[1] | 0.043060268 |
|  |  | T[2] | 0.045998142 |
|  |  | T[3] | 0.031839496 |
| Cyanobacteria | Chloroplast | T[0] | 0.020862309 |
|  |  | T[R] | 0.011650182 |
|  |  | T[1] | 0.026512447 |
|  |  | T[2] | 0.028141321 |
|  |  | T[3] | 0.018079461 |
|  | Cyanobacteriales | T[0] | 0.19073226 |
| Cyanobacteria | Cyanobacteriales | T[R] | 0.389427243 |
|  |  | T[1] | 0.333676493 |
|  |  | T[2] | 0.292463654 |
|  |  | T[3] | 0.350595706 |
|  | Unknown Oxyphotobacteria | T[R] | 0.013481119 |
|  |  | T[1] | 0.015453064 |
|  |  | T[2] | 0.013590863 |
|  |  | T[3] | 0.015066421 |
|  | Thermosynechococcales | T[0] | 0.010799313 |
| Acidobacteriota | Bryobacterales | T[R] | 0.005153526 |
| Bacteroidota | Chitinophagales | T[0] | 0.006236605 |
|  |  | T[R] | 0.005163481 |
|  |  | T[2] | 0.007262191 |
|  | Cytophagales | T[0] | 0.009163053 |
|  |  | T[R] | 0.096453172 |
|  |  | T[1] | 0.11996715 |
|  |  | T[2] | 0.079192234 |
|  |  | T[3] | 0.090668802 |
| Chloroflexi | Kallotenuales | T[R] | 0.00838268 |
|  |  | T[2] | 0.01633687 |
|  |  | T[3] | 0.008408777 |
|  | Thermomicrobiales | T[2] | 0.005146914 |
| Gemmatimonadota | Gemmatimonadales | T[R] | 0.005436359 |
|  |  | T[2] | 0.006908222 |
|  | Longimicrobiales | T[R] | 0.016006999 |
|  |  | T[2] | 0.005421611 |
| Myxococcota | Haliangiales | T[R] | 0.005678943 |
|  |  | T[3] | 0.006010924 |
|  | Myxococcales | T[R] | 0.020299082 |
|  |  | T[1] | 0.010755355 |
|  |  | T[2] | 0.013671351 |
|  |  | T[3] | 0.014621574 |
|  | Nannocystales | T[R] | 0.008809035 |
|  |  | T[1] | 0.01090915 |
|  |  | T[2] | 0.005558526 |
|  |  | T[3] | 0.006743362 |
|  | Polyangiales | T[R] | 0.008294912 |
|  |  | T[1] | 0.008084686 |
|  |  | T[2] | 0.005990088 |
|  |  | T[3] | 0.006645987 |
| Proteobacteria | Acetobacterales | T[R] | 0.014783181 |
|  |  | T[1] | 0.012028303 |
|  |  | T[2] | 0.016252769 |
|  |  | T[3] | 0.016407297 |
|  | Azospirillales | T[0] | 0.030702093 |
|  | Burkholderiales | T[0] | 0.056796051 |
| Proteobacteria | Burkholderiales | T[R] | 0.012959379 |
|  |  | T[1] | 0.010671657 |
|  |  | T[2] | 0.013591631 |
|  |  | T[3] | 0.012713217 |
|  | Caulobacterales | T[0] | 0.012899179 |
|  |  | T[R] | 0.022042144 |
|  |  | T[1] | 0.034822507 |
|  |  | T[2] | 0.020704156 |
|  |  | T[3] | 0.024863575 |
|  | Pseudomonadales | T[0] | 0.013867311 |
|  | Rhizobiales | T[0] | 0.087344626 |
|  |  | T[R] | 0.043724525 |
|  |  | T[1] | 0.063099159 |
|  |  | T[2] | 0.070129812 |
|  |  | T[3] | 0.068004988 |
|  | Rhodobacterales | T[0] | 0.00551081 |
|  |  | T[R] | 0.016305263 |
|  |  | T[1] | 0.016673137 |
|  |  | T[2] | 0.021005185 |
|  |  | T[3] | 0.016425317 |
|  | Sphingomonadales | T[0] | 0.018796014 |
|  |  | T[R] | 0.038019558 |
|  |  | T[1] | 0.044013044 |
|  |  | T[2] | 0.044435076 |
|  |  | T[3] | 0.033523863 |
| Verrucomicrobiota | Chthoniobacterales | T[R] | 0.012024879 |
|  |  | T[1] | 0.066714297 |
|  |  | T[2] | 0.044130375 |
|  |  | T[3] | 0.094877543 |

**Table A.7.** P-values of the Dunn tests between time points on the relative abundance of the actinobacterial and cyanobacterial orders. Bold numbers are significant (<0.05).

| Comparison | Cyanobacteria | Actinobacteria |
| --- | --- | --- |
| T[0] - T[1] | **6.13E-10** | **8.15E-13** |
| T[0] - T[2] | **5.13E-08** | **2.52E-14** |
| T[0] - T[3] | **5.98E-10** | **6.89E-11** |
| T[0] - T[R] | **6.52E-10** | **6.47E-14** |
| T[1] - T[3] | 0.498128567 | 0.228324880 |
| T[1] - T[2] | 0.191797281 | 0.294405229 |
| T[1] - T[R] | 0.495469243 | 0.345672443 |
| T[2] - T[R] | 0.194912395 | 0.442935801 |
| T[3] - T[R] | 0.493597980 | 0.126850627 |
| T[2] - T[3] | 0.190519550 | 0.099407424 |

**Table A.8.** Abundance (in copy number (CN)) of each time points within each group of gene.

| Gene Group | Time points | Abundance |
| --- | --- | --- |
| DNA conservation | T[0] | 20590 |
|  | T[R] | 91433 |
|  | T[1] | 92496 |
|  | T[2] | 78321 |
|  | T[3] | 81983 |
| DNA repair and degradation | T[0] | 13579 |
|  | T[R] | 66132 |
|  | T[1] | 67048 |
|  | T[2] | 55948 |
|  | T[3] | 58457 |
| Light energy or sensing | T[0] | 85 |
|  | T[R] | 43 |
|  | T[1] | 59 |
|  | T[2] | 64 |
|  | T[3] | 17 |
| Lithotrophs | T[0] | 7554 |
|  | T[R] | 37972 |
|  | T[1] | 38632 |
|  | T[2] | 31341 |
|  | T[3] | 32758 |
| Nitrogen | T[0] | 10027 |
|  | T[R] | 50708 |
|  | T[1] | 58068 |
|  | T[2] | 48225 |
|  | T[3] | 45638 |
| Organotrophs | T[0] | 60007 |
|  | T[R] | 108275 |
|  | T[1] | 111044 |
|  | T[2] | 88557 |
|  | T[3] | 89148 |
| Phototrophy | T[0] | 50301 |
|  | T[R] | 445432 |
|  | T[1] | 425819 |
|  | T[2] | 342188 |
|  | T[3] | 407532 |
| ROS-damage prevention | T[0] | 26126 |
|  | T[R] | 138367 |
|  | T[1] | 143726 |
|  | T[2] | 121507 |
|  | T[3] | 126677 |
| Sensing & motility | T[0] | 31075 |
|  | T[R] | 81947 |
|  | T[1] | 92070 |
| Sensing & motility | T[2] | 90844 |
|  | T[3] | 74436 |
| Sporulation capsule & C-storage | T[0] | 7302 |
|  | T[R] | 48141 |
|  | T[1] | 48944 |
|  | T[2] | 38998 |
|  | T[3] | 42341 |

**Table A.9.** Chi-square values and p-values of the Dunn tests between time points done on the functional prediction results. Bold numbers are significant (< 0.05)

| Comparison | DNA Conservation | DNA Repair | Light energy | Lithotrophy | Nitrogen |
| --- | --- | --- | --- | --- | --- |
| T[0] - T[1] | **0.00489293** | **0.00293591** | 0.19388801 | **0.00030834** | **4.22E-17** |
| T[0] - T[2] | **0.02033475** | **0.0097339** | 0.44831059 | **0.00749298** | **8.12E-12** |
| T[0] - T[3] | **0.02906312** | **0.01558465** | 0.2755365 | **0.01520981** | **8.15E-10** |
| T[0] - T[R] | **0.02863121** | **0.00520412** | 0.37990911 | **0.00162909** | **1.30E-12** |
| T[1] - T[3] | 0.21328302 | 0.24422913 | **0.04593218** | 0.072911 | **0.00403999** |
| T[1] - T[2] | 0.26782432 | 0.31450078 | 0.19843215 | 0.12613918 | **0.03331074** |
| T[1] - T[R] | 0.21549186 | 0.41187192 | 0.25970621 | 0.28894927 | 0.06264837 |
| T[2] - T[3] | 0.43027457 | 0.41697806 | 0.20090041 | 0.37842665 | 0.20768207 |
| T[2] - T[R] | 0.43325114 | 0.3972775 | 0.41957403 | 0.27813996 | 0.38159137 |
| T[3] - T[R] | 0.49697916 | 0.31916653 | 0.14884666 | 0.184595 | 0.13225512 |
| **Chi-square** | 7.021623874 | 8.913054423 | 2.944440598 | 12.94456096 | 75.97487673 |

| Comparisons | Organotrophy | Phototrophy | ROS | Motility | Sporulation |
| --- | --- | --- | --- | --- | --- |
| T[0] - T[1] | **0.00189691** | **5.38E-36** | **5.70E-06** | **1.09E-18** | **7.48E-05** |
| T[0] - T[2] | **0.02110913** | **8.46E-17** | **2.77E-05** | **2.10E-14** | **3.01E-09** |
| T[0] - T[3] | **0.04112206** | **8.77E-26** | **8.96E-05** | **2.41E-11** | **0.0008838** |
| T[0] - T[R] | **0.01948721** | **4.27E-35** | **6.29E-05** | **1.22E-14** | **0.00040669** |
| T[1] - T[3] | 0.09077588 | **0.00930641** | 0.22920864 | **0.00608522** | 0.22132375 |
| T[1] - T[2] | 0.15937532 | **5.23E-07** | 0.33993514 | 0.08422197 | **0.00969216** |
| T[1] - T[R] | 0.16882618 | 0.42394382 | 0.26107583 | 0.09739203 | 0.30432515 |
| T[2] - T[3] | 0.3673185 | **0.00570786** | 0.37114762 | 0.12925022 | **0.00094887** |
| T[2] - T[R] | 0.48475837 | **1.36E-06** | 0.41005945 | 0.46785414 | **0.00218557** |
| T[3] - T[R] | 0.35302059 | **0.01533294** | 0.45960721 | 0.11302291 | 0.39907998 |
| **Chi-square** | 8.60329063 | 195.4178784 | 22.91571278 | 87.21260295 | 34.5684137 |

**Figure A.1.**
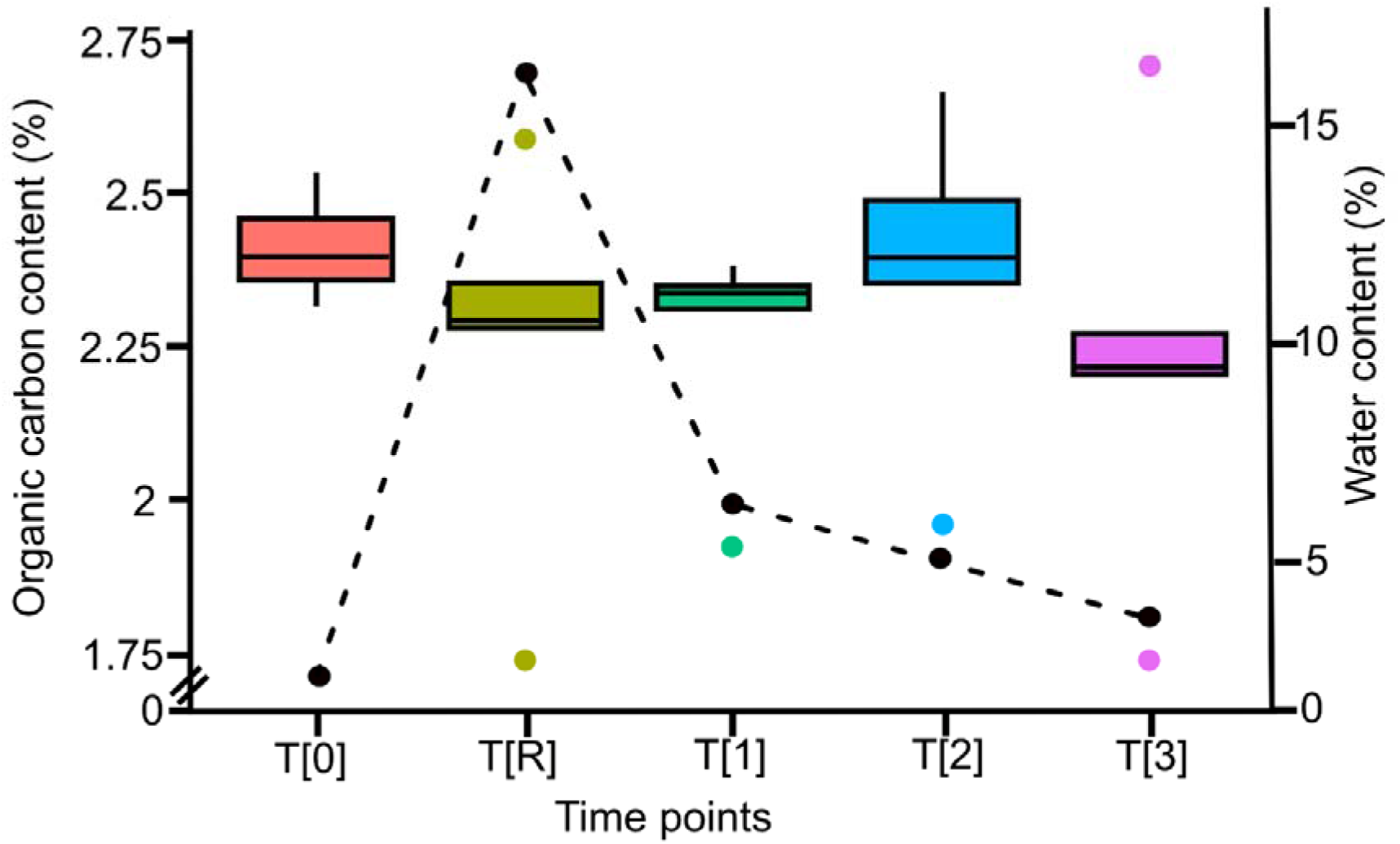
Barplot of the organic carbon content for each sampling point.

**Figure A.2.**
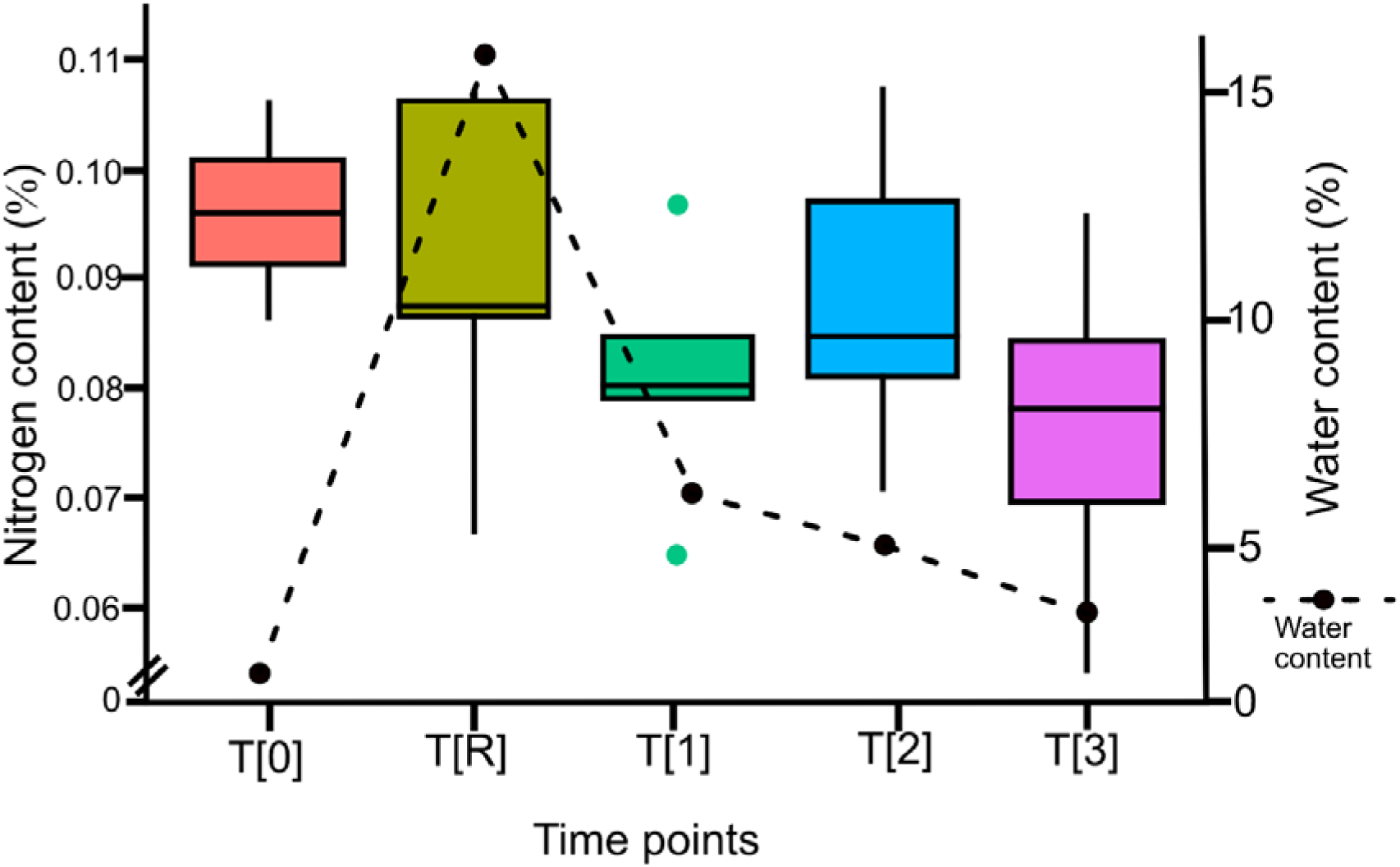
Barplot of the nitrogen content (in g) for each time point.

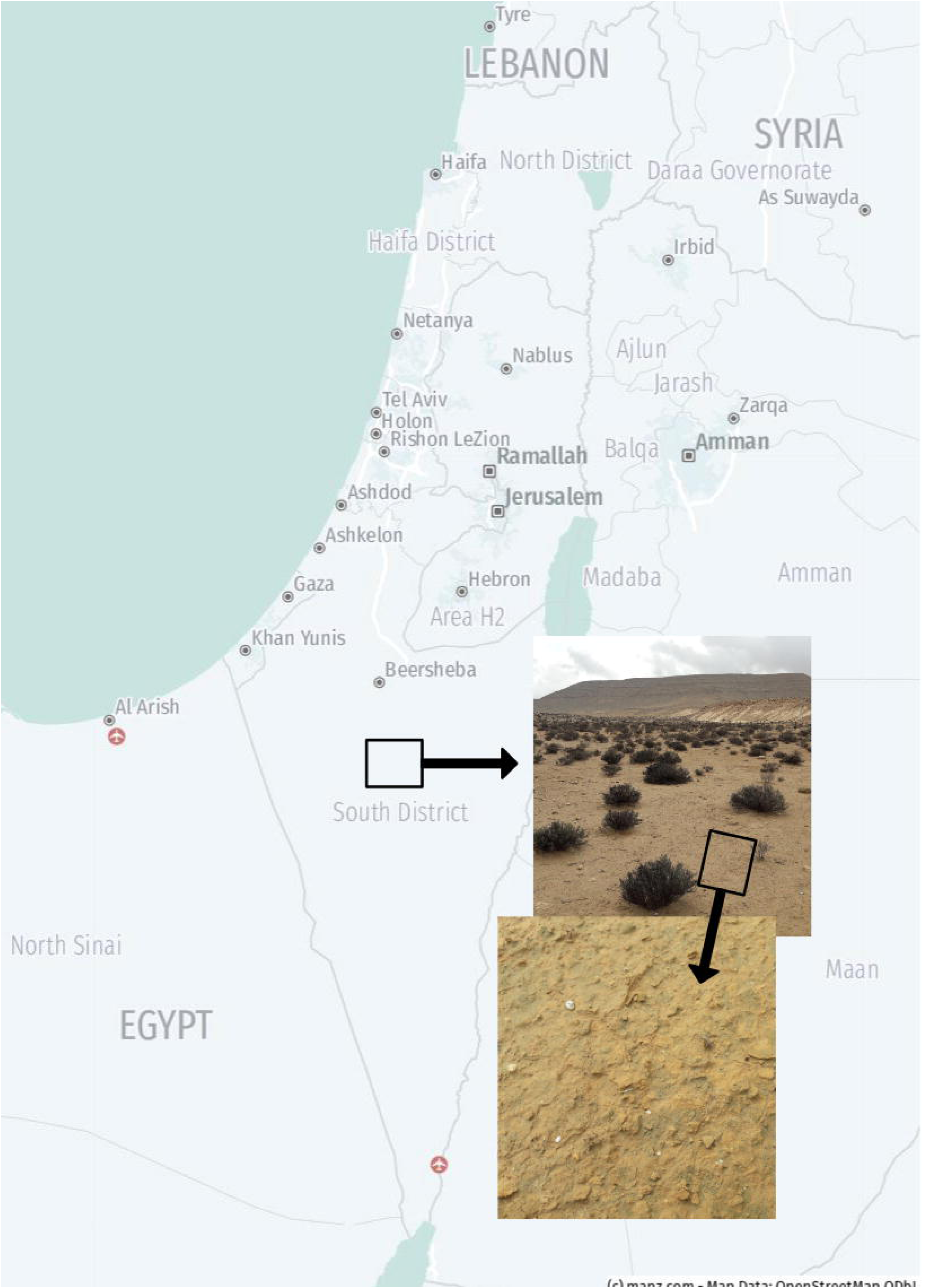

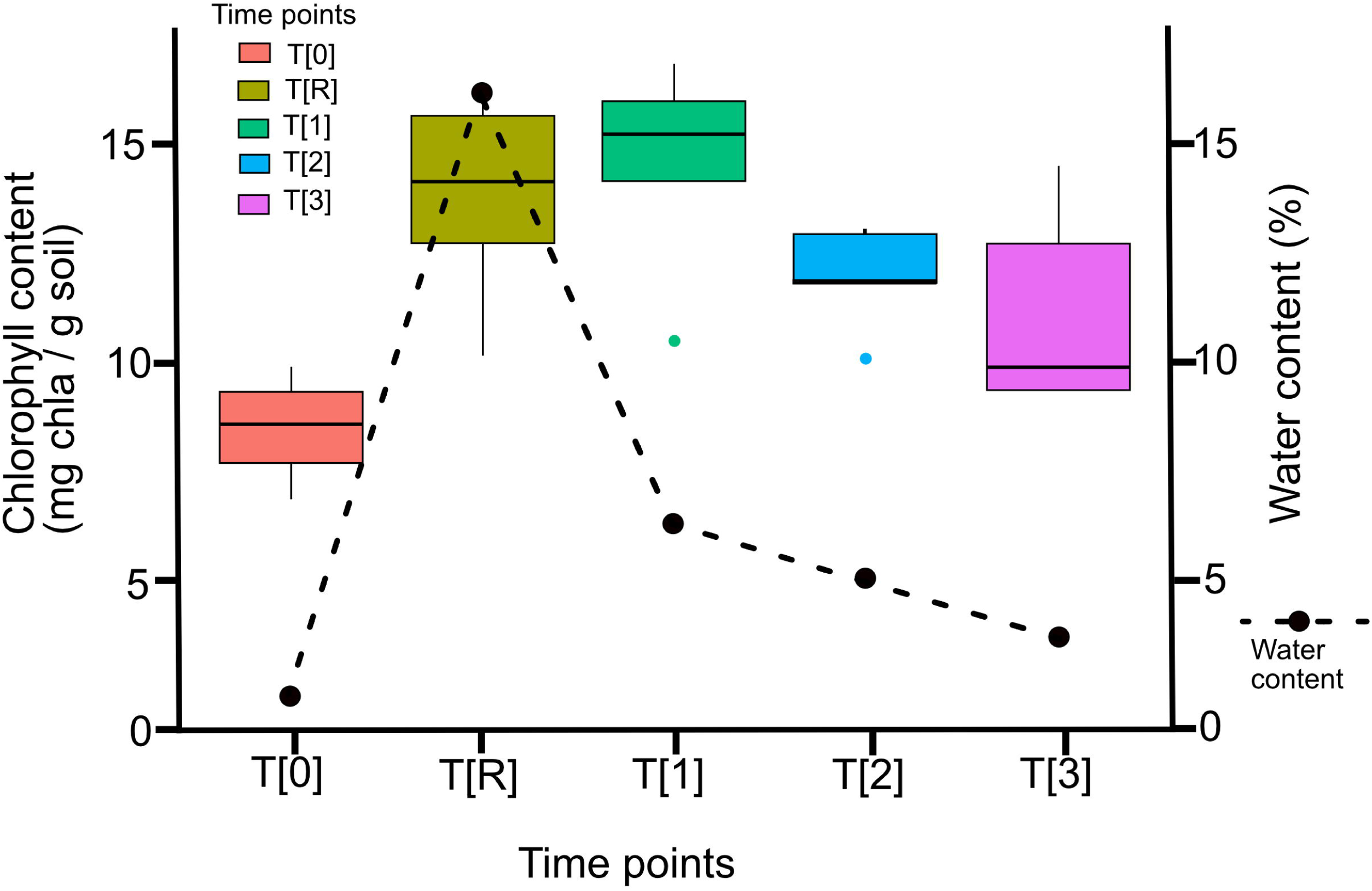

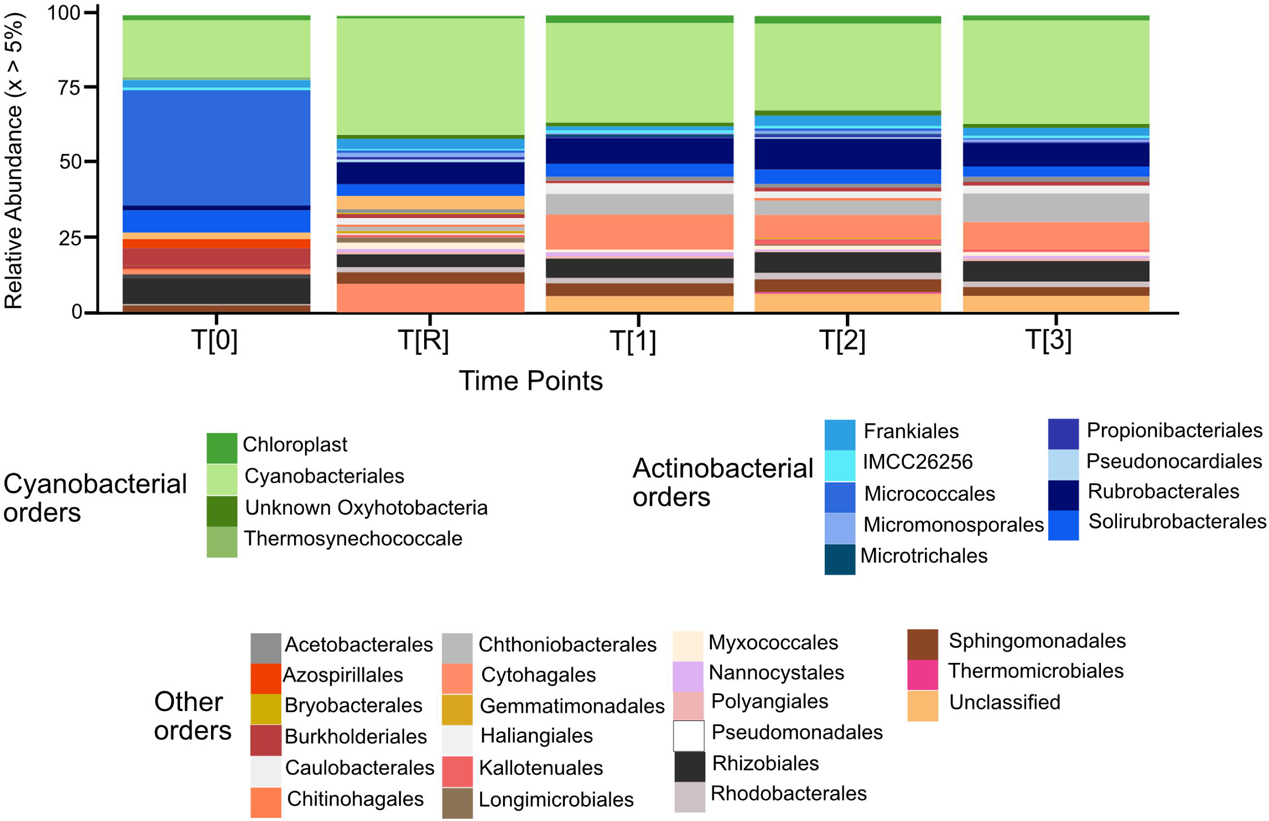

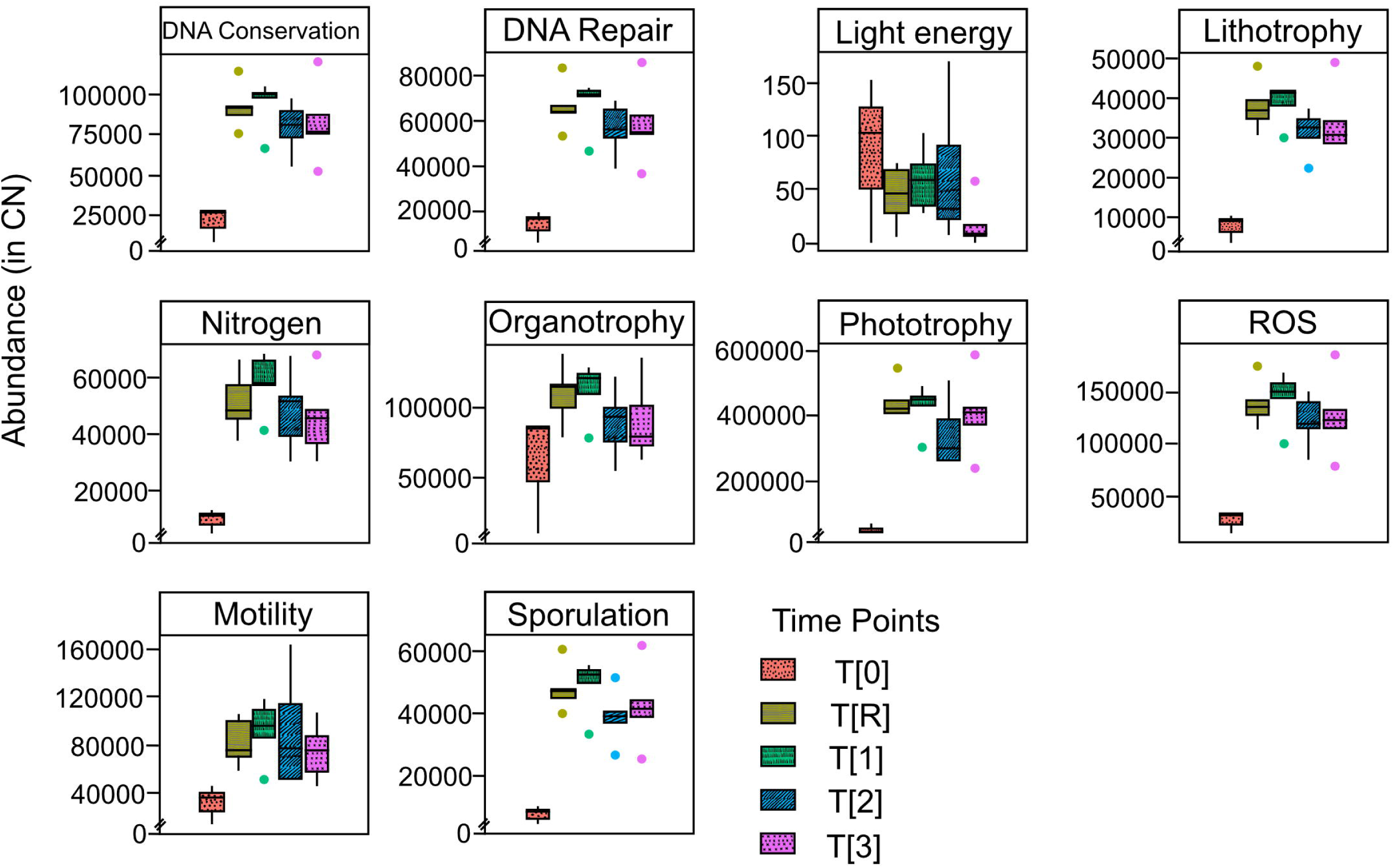

